# Impact of Partition Models on Phylogenetic Inference and Divergence Times of Lampyridae from Mitochondrial Genomes

**DOI:** 10.1101/2025.08.19.671050

**Authors:** Sebastian Höhna, Haoqing Du, Ana Catalán

## Abstract

Mitochondrial genomes are frequently used for phylogenetic inference due to their availability and cost-efficient sequencing. In most mitogenomic phylogenetic analyses, only the two ribosomal RNA and 13 protein coding genes are used. For such multi-locus datasets it has long been established that appropriate data partition models, e.g., partitioning by gene type and/or codon position, are necessary for robust phylogenetic inference. While most studies focused on the impact of partition models on the tree topology, little is known about the impact on divergence time estimation. Furthermore, although modeling among site rate variation within a partition is common practice, the extent of substitution rate variation among partitions is less explored.

Here we study the impact of four partition model strategies: (i) no subdivision of the data (*uniform*), (ii) partitioning by *gene*, (iii) partitioning by *codon* position, and (iv) partitioning by both gene and codon position (*combined*). We explore the impact of partition models on divergence time estimation in fireflies (Coleoptera: Lampyridae). To this end, we sequenced mitochondrial genomes from 22 firefly species from Europe and Central America and combined these with 82 published mitochondrial genomes. Our results represent the most comprehensive time-calibrated phylogeny of fireflies to day. While the partition models had an impact on the inferred tree topology, the divergence times were almost not affected. Furthermore, we observed up to 3-fold substitution rate variation across genes and additionally more than to 10-fold substitution rate variation across codon positions. Our study gives insights into best practices of performing partitioned-data time-calibrated phylogenetic inference from mitochondrial genomes and multi-locus datasets in general.

## 1 Introduction

Multi-locus datasets consist of loci with possibly different characteristics (e.g., non-coding regions, protein coding genes and ribosomal RNA) and evolutionary constraints (e.g., faster/slower rate of evolution or different base compositions). In phylogenetics, it is therefore common practice to partition multi-locus datasets (Yang, 1996; Duchêne et al., 2011; Cameron, 2014; Nascimento et al., 2017). The standard approach is to divide the full dataset by shared features: by locus type (e.g., coding vs. non-coding), by locus (e.g., each locus separately), by codon position, or any combination thereof (Kainer and Lanfear, 2015; Timmermans et al., 2016; Höhna et al., 2017). Each data subset within the overall dataset can receive its own set of substitution model parameters, or parameters can be linked across data subsets. Some tools exist to identify and assign candidate substitution models to data subsets and to merge data subsets together (Lanfear et al., 2012, 2014; Kalyaanamoorthy et al., 2017). In general, wrong parameter assumptions of shared/linked substitution process across data subsets can lead to biased topology and branch length estimates (Marshall et al., 2006).

The primary and most common focus of partitioned data phylogenetic models lies on the substitution model and its parameters (the exchangeability rates, stationary frequencies, and among site rate variation) (Duchêne et al., 2011; Timmermans et al., 2016; Yang et al., 2018). Interestingly, much less focus has been laid on modelling variation in overall substitution rates among data subsets, i.e., modelling among partition rate variation. Inappropriate modelling of among partition rate variation can strongly affect branch-lengths (Marshall et al., 2006) but the choice of data partitioning on divergence time estimation is more intricate (Angelis et al., 2018). Currently, the standard approach to model among partition rate variation is to apply a Gamma-Dirichlet prior distribution, i.e., a Gamma hyperprior distribution on the overall substitution rate and a Dirichlet distribution for the relative rates among partitions (Dos Reis et al., 2014; Höhna et al., 2017). That is, the Dirichlet distribution assigns *a priori* equal among partition rates and if a flat Dirichlet distribution is used, then any among partition rate variation is equally probable *a priori*. Thus, an among partition rate variation of several orders of magnitude is equally likely as equal per-partition rates. Empirical observations show that different types of genes evolve at different speeds, and particularly within protein coding genes the third codon positions evolve much faster than the first and second codon position (Shapiro et al., 2006). However, extreme among partition rate variation of several orders of magnitude, as postulated by the flat Dirichlet prior distribution, is biologically unrealistic. The question remains how much among partition rate variation exists in empirical data sets to inspire better models of among partition rate variation.

In this study, we focus on the impact of partition model choice on divergence time estimation using mitochondrial genomes in fireflies (Coleoptera: Lampyridae). Mitochondrial genomes are very popular for phylogenomics especially in insects (Cameron, 2014) and are well suited to explore the impact of among partition rate variation on phylogeny (i.e., tree topology) and divergence time estimation (Duchêne et al., 2011; Timmermans et al., 2016; Yang et al., 2018). These mitochondrial genome “phylogenomic” studies often use the 13 mitochondrial protein coding genes as well as 2 ribosomal RNA genes. Other data sources, such as tRNAs, are commonly omitted for phylogenomic inference. Given their molecular structure, it is expected that rRNA genes evolve at a different speed compared with protein coding genes. Thus, the maximal number of data subsets is 41 (dividing the 13 protein coding genes by codon position plus the two rRNA genes) making the exploration computationally feasible while being biologically relevant. Additionally, mitochondrial loci can be considered to evolve along the same phylogeny due to lack of recombination and therefore loci can be concatenated without concern about gene tree-species tree discordance. Finally, mitochondrial genomes remain a popular choice for phylogenomic inference due to cost-efficient sequencing.

Our study system here are fireflies, beetles well-known due to their bioluminescence and comprising over 2,500 extant species (McDermott, 1964; Branham and Wenzel, 2001; Stanger-Hall et al., 2007; Martin et al., 2017, 2019; Ferreira et al., 2020). Despite their charismatic features and popularity, phylogenetic studies based on molecular data have covered little of their diversity. Recent divergence time estimates indicate that crown fireflies originated up to 140 million years ago (Powell et al., 2022; Höhna et al., 2025) with the oldest known fossil from Burmese amber being 99 million years old (Kazantsev, 2015; Haug et al., 2023; Cai et al., 2024). Despite these recent efforts (Martin et al., 2019; Powell et al., 2022; Höhna et al., 2025), the phylogenetic relation among many firefly species, genera and especially subfamilies is still unknown and their divergence times need further corroboration. Additionally, there is strong under-representation of genetic material from Neotropical (Catalán et al., 2025b) and European fireflies, while more efforts have been undertaken in North America and Asia, causing biased sampling and imbalanced phylogenetic relationships among fireflies.

Here, we sequenced 22 new mitochondrial genomes of fireflies and combined these with 82 already published firefly mitochondrial genomes. We explored four different data partition strategies: (1) *uniform* (i.e., no partition) where all genes are concatenated and assumed to evolve under the same substitution model, (2) by *gene* where each genes evolves under its own substitution model (15 data subsets), (3) by *codon* position where all first, second and third codon position of the protein coding genes are merged into three separate data subsets, each assumed to evolve under its own substitution model (4 data subsets), and (4) *combined* by gene and codon position where the first, second and third codon positions of each gene are placed into a separate data subset and assumed to evolve under their own substitution model (41 data subsets). For each data subset, we assumed that molecular sequences evolve along the same phylogeny but with different overall substitution rates (among partition rate variation). We performed phylogenetic inferences of single genes and concatenations of all genes for unrooted phylogenies to investigate the correlation of single gene tree lengths and among partition rate variation in concatenated analyses. Furthermore, we performed phylogenetic inferences using unrooted and time-calibrates phylogenetic models to investigate the connection between substitution rates, clock rates, and divergence times. We time-calibrated the phylogeny using the fossilized birth-death process (Heath et al., 2014) as precise node calibrations are lacking (Parham et al., 2012). Divergence times were inferred in `RevBayes` (Höhna et al., 2016) under a relaxed-clock mixture model (Darlim and Hoehna, 2024). Our results show that partition models impact both topology and substitution rate estimates, however, divergence times estimates are almost not affected because substitution rate variation was absorved by variation in clock rates.

## 2 Methods

### 2.1 Data Sampling and Sequencing

We collected 22 firefly species from Europe and Central America (Figure S1 and Table S1). Bioluminiscent adults (*Luciola, Photinus, Photuris* and *Bicellonycha*) were collected using an insect net and dark adults were collected using light traps (*Lamprohiza* and *Lampyris*). High molecular weight DNA was extracted from single males using the MagAttract HMW DNA kit (Qiagen) following manufacturer’s guidelines. DNA fragment sizes and integrity were checked with a 1% agarose gel. Long DNA fragments were sequenced from a single male individual using Nanopore PromethION sequencer. Flongle flow cells were run for 72 hours. Nanopore sequencing was done by the Gene Center at the LMU München, Munich, Germany and base calling was done using Oxford Nanopore’s Basecaller `dorado` (https://github.com/nanoporetech/dorado).

### 2.2 Mitochondrial Genome Assembly

Standard pipelines for mitochondrial genome assembly expect short-read Illumina DNA sequences but can be used with long-read Nanopore sequencing data having only 1% error rate. Thus, we explored two bioinformatic pipelines, using Nanopore reads, for mitochondrial genome assembly: `MitoFinder` (Allio et al., 2020) `and MitoHifi` (Uliano-Silva et al., 2023). Each pipeline was run with four reference mitochondrial genomes as our best guess for most closely related sequenced species (*Lampyris noctiluca, Photinus signaticolis* (Koken and Gastineau, 2024), *Bicellonycha lividipennis* (Amaral et al., 2016), *Curtos bilineatus* (Chen et al., 2019)). Then, mitochondrial genomes were annotated using `Mitoz v3.6` (Meng et al., 2019). From the 8 different mitochondrial genome assembly settings, we selected the best assembly according to the following criteria: (1) all 13 protein coding genes and 2 rRNA genes were found (completeness), (2) the shortest mitochondrial genome assembly, (3) choosing the hit to the phylogenetically closest reference genome.

### 2.3 Sequence Alignments

We downloaded 82 previously published firefly mitochondrial genomes (see Table S1; (Bae et al., 2004; Amaral et al., 2016; Mu et al., 2016; Fan and Fu, 2017; Wang et al., 2017; Fallon et al., 2018; Hu and Fu, 2018b,a; Chen et al., 2019; Sriboonlert and Wonnapinij, 2019; Yang and Fu, 2019; Zhang and Fu, 2019; Kim et al., 2020; Liu and Fu, 2020; Wang et al., 2021; Ge et al., 2022; Ji and Xu, 2022; Koken and Gastineau, 2024)). We combined these published genomes with our 22 mitochondrial genomes. We aligned the 13 protein coding genes using `MACSE v2` (Ranwez et al., 2018), using the insect mitochondrial code. We also used `MACSE v2` to replace artificial stop codons with ‘NNN’ if these occurred within a protein coding sequence. Furthermore, we trimmed the alignments using `trimAl` (Capella-Gutiérrez et al., 2009) with the option `automated1`. For the two ribosomal RNA genes 12S and 16S, we used the alignment software `MAFFT v7` (Katoh and Standley, 2013) and `trimAl` (Capella-Gutiérrez et al., 2009) with the option `automated1`. Alignments of these 15 loci were used as input into `RevBayes` (Höhna et al., 2016), where the loci were subset and concatenated according to the data partition strategies.

### 2.4 Phylogenetic inference of individual genes

We performed phylogenetic inference of individual gene alignments to identify outlier sequences/taxa and to explore the among-gene overall substitution rate variation. Our individual gene phylogenetic analyses were performed using unrooted phylogenies where the branch lengths are in units of expected substitutions per site. Thus, the total tree length corresponds to the overall expected number of substitutions per site for this gene, and tree length estimates can be compared between genes to assess the among-gene substitution rate variation.

We explored two partition strategies for these single gene analyses: no partition (i.e., uniform) and partitioning by codon position (only applied to the protein coding genes and not the rRNA genes). Nucleotide sequence evolution was modelled within each data subset by an independent GTR+Γ+I substitution model (Tavaré, 1986; Yang, 1994). We assumed flat Dirichlet prior distributions for the stationary frequencies and exchangeability rates and a uniform prior distribution on the α parameter between 0 and 10^8^ for the among-site rate variation (Fabreti and Höhna, 2023). The proportion of invariant sites had a flat Beta(1,1) prior distribution. Furthermore, we assumed equal prior probabilities for each tree topology and an exponential distribution with mean 0.1 for each branch length. We ran 4 independent Markov chain Monte Carlo (MCMC) replicates for 100,000 iterations using 893.5 moves per iteration in `RevBayes`(Höhna et al., 2016). We tested for failure to converge using the `R` package `convenience`(Fabreti and Höhna, 2022).

### 2.5 Phylogenetic inference of mitochondrial genomes with different partition models

We performed species-tree phylogenetic inference from the mitochondrial genomes of fireflies under different partition models in the Bayesian phylogenetics software `RevBayes` (Höhna et al., 2016). Specifically, we divided the full dataset (13 protein coding genes and 2 rRNA genes) using four different partition strategies: (i) *uniform*, i.e., all sites in the alignment evolve under the same substitution process with the same shared parameters, (ii) by *gene*, i.e., the alignment is split into 15 subsets according to the genes, (iii) by *codon* position, i.e., four data subsets where the two rRNA genes are merged, all 1st codon positions of the protein coding genes are merged together, all 2nd codon positions are merged, and all 3rd codon positions are merged, and (iv) by gene and codon position *combined*, i.e., the two rRNA genes are split into separate data subsets and each protein coding gene is split into three data subsets based on codon position (41 data subsets in total). We used the same prior distributions on the substitution model parameters as in our individual gene phylogenetic analyses (see above).

When we applied a partition model for the data subsets, i.e., all analyses except the uniform partition, then we applied a flat Dirichlet prior distribution on among partition rate variation. The per-partition rate multipliers were rescaled to an average rate of 1.0, such that branch lengths are units expected number of substitutions for the average data subsets. Moreover, we assumed an independent exponential prior distribution with a mean of 0.1 for each branch, thus an *a priori* expected tree length of 0.1 × (2*n* − 3) = 20.5. While these analyses focused primarily on the inference of the tree topology, they also established a baseline of the impact of the partition models on the overall tree length, which is indicative for the clock rate in a divergence time analysis.

We ran four MCMC replicates for 100,000 iterations using 1351.5 moves per iteration in the parallel version `RevBayes`(Höhna et al., 2016; Smith et al., 2024). We tested for failure to converge using `the R` package `convenience`(Fabreti and Höhna, 2022).

### 2.6 Time calibrated phylogenetic inference

We performed joint phylogenetic inference and divergence time estimation in the Bayesian phylogenetics software `RevBayes` (Höhna et al., 2016). The aim of this analysis was two-fold: first, we aimed to infer a time-calibrated phylogeny of fireflies, and second, we aimed to explore the impact of partition models on divergence time estimates. Thus, we used the same four partition models as in the unrooted phylogeny analyses above.

We identified three firefly fossils to time-calibrate our phylogenetic analyses: †*Lampyris orciluca*(Heer, 1865), †*L*amprohiza fossilis(Kazantsev, 2012) and †*Photinus kazantsevi*(Alekseev, 2019) (see Table 1). We did not use †*Lucidota prima*(Wickham, 1912) because of lack of sequenced taxa to calibrate a node (there is currently no available mitochondrial genome for *Lucidota*) and †*Electrotreta rasnitsyni*(Kazantsev, 2012) due to uncertain phylogenetic placement (see (Powell et al., 2022; Höhna et al., 2025)). Furthermore, we did not use †*Protoluciola albertalleni*(Kazantsev, 2015) and †*Flammarionella hehaikuni*(Cai et al., 2024), two Burmese amber fossils of age 99Ma assigned to the subfamily Luciolinae, but instead used these as minimum calibrations for the root age of all Lampyridae. Thus, we assigned a broad uniform prior distribution between 99 and 200 MYA on the crown of fireflies.

**Table 1.** Fossils and calibration constraints for the time-calibrated divergence time analysis.

| Fossil taxon | Age range (MA) | Reference | Monophyletic clade constraint |
| --- | --- | --- | --- |
| †*Lampyrus orciluca* | 11.608 – 12.7 | (Heer, 1865) | †*Lampyrus orciluca*, *Lampyrus noctiluca* sp1, *Lampyrus noctiluca* sp2, *Lampyrus noctiluca* sui, *Lampyrus zenkeri*, *Lampyrus iberica*, *Lampyrus lareynii*, *Nyctophila reichii* |
| †*Lamprohiza fossilis* | 23.03 – 28.4 | (Kazantsev, 2012) | †*Lamprohiza fossilis*, *Lamprohiza mulsantii*, *Lamprohiza paulinoi*, *Lamprohiza germari*, *Lamprohiza splendidula* |
| †*Photinus kazantsevi* | 33.9 – 37.2 | (Alekseev, 2019) | †*Photinus kazantsevi*, *Photinus* sp1, *Photinus* sp2, *Photinus* sp3, *Photinus* sp4, *Photinus schusterii*, *Photinus pyralis*, *Ellychnia corrusca*, *Photinus signaticollis* sp1, *Photinus signaticollis* sp2 |

The fossil record of fireflies is extremely sparse and their phylogenetic position comparably uncertain, particularly concerning whether these fossil are stem or crown fossils, which complicates traditional phylogenetic node calibrations. Therefore, we used instead the fossilized birth-death process (Heath et al., 2014) which uses the fossils as tips in the phylogeny and jointly applies a prior distribution on the divergence times and fossil sampling ages. Furthermore, we applied fossil-age uncertainties using uniform distribution between specified minimum and maximum ages (see Table 1) to reflect uncertainty in the ages of the fossils themselves (Barido-Sottani et al., 2019).

We performed our time-calibrated phylogenetic inference under a relaxed-clock mixture model (Darlim and Hoehna, 2024) composed of an uncorrelated exponential (Drummond et al., 2006), independent gamma (Lepage et al., 2007) and uncorrelated lognormal prior distribution on branch-specific clock rates (Drummond et al., 2006). We chose a uniform prior distribution between 0 and 10 substitutions per million years per site on the mean clock rate, and assigned exponential prior distributions on the variance parameter of the independent gamma and uncorrelated lognormal prior models. We ran 4 independent Markov chain Monte Carlo (MCMC) replicates for 100,000 iterations using 1351.5 moves per iteration in the parallel version `RevBayes`(Höhna et al., 2016; Smith et al., 2024). We tested for failure to converge using `the R` package `convenience`(Fabreti and Höhna, 2022).

## 3 Results

### 3.1 Twenty-two new Lampyridae mitochondrial genomes

We sequenced and assembled 22 new mitochondrial genomes of fireflies, including several new and yet described firefly species from Guatemala (Fig. 1). In general, we found that `MitoHifi` (Uliano-Silva et al., 2023) worked better than `MitoFinder` (Allio et al., 2020) for our long-read sequences, likely due to the higher error rate of the Nanopore PromethION sequencer, when compared to Illumina sequencing.

**Figure 1.**
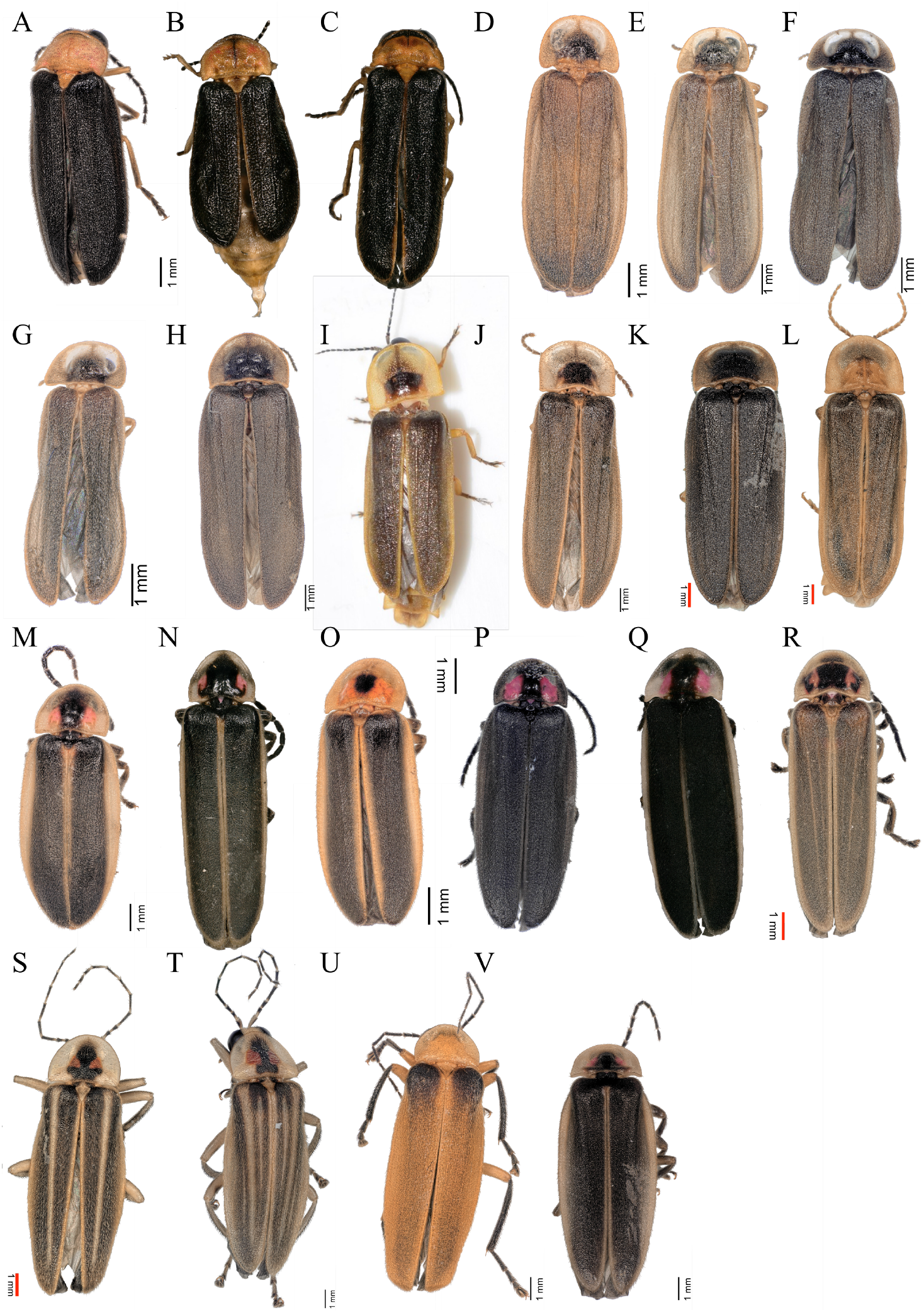
Staked images of firefly dorsals whose mitochondrial genomes were newly assembled. Mostly males were imaged unless indicated. **(A)** Luciola novaki, **(B)** Luciola lusitanica(female), **(C)** Luciola italica, **(D)** Lamprohiza paulinoi, **(E)** Lamprohiza mulsanti, **(F)** Lamprohiza splendidula, **(G)** Lamprohiza germari, **(H)** Lampyris noctiluca, **(I)** Lampyris zenkeri, **(J)** Lampyris lareynii, **(K)** Lampyris iberica, **(L)** Nyctophila reichii, **(M)** Photinus sp1, **(N)** Photinus schusteri, **(O)** Photinus sp4, **(P)** Photinus sp3, **(Q)** Photinus sp2, **(R)** Photinus signaticollis, **(S)** Photuris sp1, **(T)** Photuris sp2, **(U)** Photuris sp3 (female), **(V)** Bicellonycha sp1.

The specific choice of reference mitochondrial genome mattered, although all four chosen references were from the firefly family *Lampyridae*, but primarily for annotation. Manual inspection and curration was necessary to remove wrongly inserted sequences (e.g., duplications) and wrong single nucleotide insertions/deletions causing frame shifts. Manual curation was feasible for these mitochondrial genomes because of their relatively small size (15–18 kilobases) but was necessary (Fig. S1).

Our curated mitochondrial genomes of the 22 new fireflies contained all 13 protein coding genes, both ribosomal RNA genes, and all 22 transfer RNA (tRNA) genes. The overall lengths of the mitochondrial genomes ranged between 16.1 and 18.2 kilobases, placing our mitochondrial genomes slightly towards the larger half of published firefly mitochondrial genomes (Fig. S1). Our new sequences include the first mitochondrial genomes of the European genera *Lamprohiza* and *Nyctophila* and adds several new mitochondrial genomes to the European genus *Lampyris* and the American genera *Photuris, Photinus* and *Bicellonycha*.

### 3.2 Substitution rate variation in mitochondrial loci

We first explored the substitution rate variation of mitochondrial loci by estimating unrooted gene trees. Thus, the overall substitution rate was obtained from the gene tree length as branches are measured in expected number of substitutions and therefore the tree length is directly proportional to the substitution rate. We observed that the two rRNA genes had significantly lower substitution rates, roughly half the number as in protein coding genes (Fig. 2). When we assumed a uniform partition within each gene, then we recovered some substitution rate variation across protein coding genes (Fig. 2 top left). However, when we assumed a partition by codon position, then the variation in substitution rates among protein coding genes almost completely disappeared (Fig. 2 top right). Interestingly, the estimated tree length was very close to the prior distribution on the tree length, indicating that the by *codon* partition scheme induces more prior sensitivity on the tree length.

**Figure 2.**
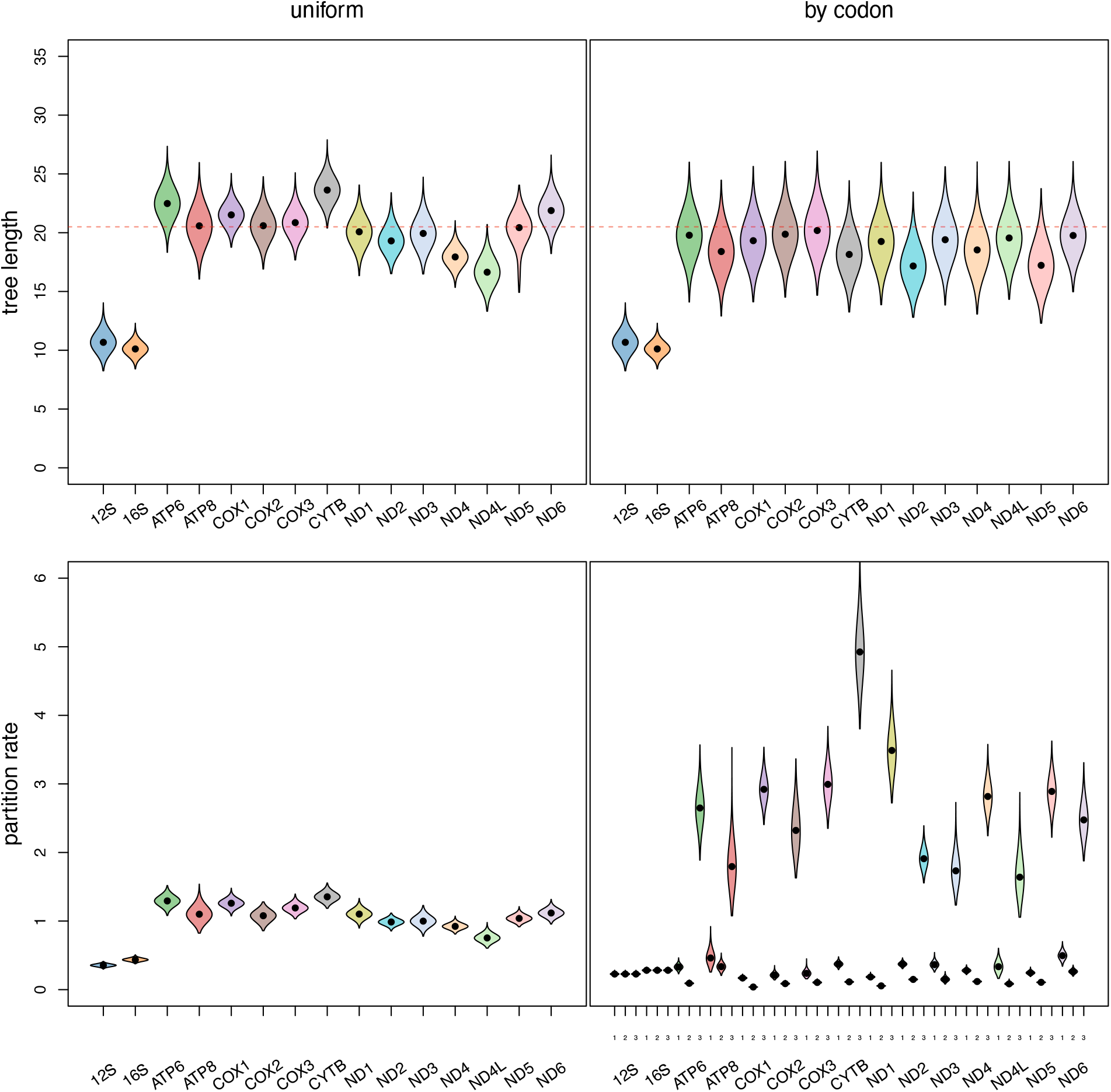
Exploration of substitution rate variation among mitochondrial loci in fireflies. We performed Bayesian inference of gene trees (top) separately and Bayesian inference of the species tree from concatenated loci (bottom) using the two rRNA (12S and 16S) and the 13 protein coding genes. Top left: Posterior distribution of tree lengths for the 15 mitochondrial genes assuming a uniform partition model, i.e., the substitution process is assumed to be homogeneous across the entire gene. The red dashed line shows the prior mean of the tree length. Top right: Posterior distribution of tree length for the 15 mitochondrial genes assuming a partition model by codon position, i.e., each protein coding gene is split into first, second and third codon position. Each of the three data subsets is modeled by its own independent substitution process but along a shared phylogeny. Note that the two rRNA genes were not split by codon position and therefore received a uniform partition. Bottom left: Posterior distribution of partition rates for each gene in a concatenated analysis when assuming the data is partition by gene but homogeneous/uniform within each gene. Bottom right: Posterior distribution of partition rates for each gene and codon position in a concatenated analysis when assuming the data is partitioned by gene and codon position (combined). Note that the two rRNA genes were not partitioned by codon, but for simplicity of visualization, we show also for the rRNA genes three partition rates (but exactly equal ones).

The partition model analyses of the concatenated genes matches the single gene phylogenetic analyses. First, when we assumed a partitioning by *gene*, then the partition rates match the observed patterns from the tree length of the single gene analyses. For example, the two rRNA genes had the shortest tree length (Fig. 2 top left) and also received the lowest partition rate (Fig. 2 bottom left). Second, when we assumed a partitioning by gene and codon position (*combined*), then we observed significant variation in partition rates among codon positions, with third codon positions receiving the highest rates and the second codon positions the lowest rates. Interestingly, we observed some rate variation between the genes at the third codon position, indicating that a combined partitioning strategy could be most appropriate. Finally, we also observed that the tree length estimates between all four partitioning strategies were different, indicating that the partitioning strategies can have an impact on divergence time estimation (Fig. 3 left).

**Figure 3.**
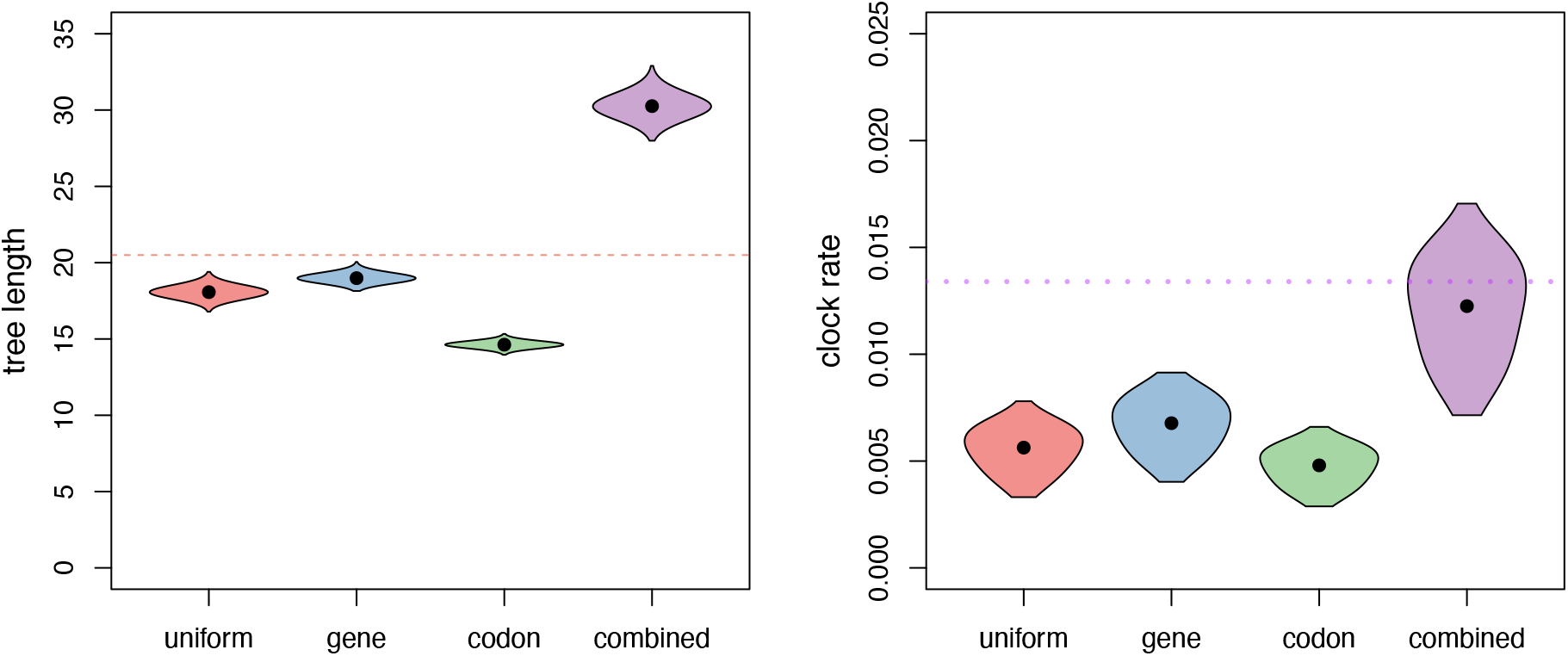
Exploration of impact of partition model on substitution and clock rates. We performed Bayesian inference of phylogeny using concatenated alignments of two rRNA and 13 protein coding genes. We performed two sets of each four analyses, the first set using an unrooted phylogeny to estimate the total tree length as a proxy of substitution rates (left), and the second set using a time-calibrated phylogeny under a relaxed-clock model to estimate the clock rates (right). Each set contained four different partition models (see description in the main text): uniform, gene, codon, and combined. The red dashed line on the left shows the prior mean on the three length, and the dotted purple line on the right shows the previously published mitochondrial clock rate in coleoptera (Pons et al., 2010).

The data partition strategy also had an impact on the inferred gene tree topologies (Figs. S2–S14), the posterior probabilities of selected clades (Figs. S15 & S16) and species tree topology (Fig. 4 and S17–S20). For the concatenated alignment species tree inferences where we explored four different partition models, we observed the strongest difference on posterior probabilities of species relationships between the *uniform* partition model and all other partition models (Fig. 4 and S21). Furthermore, we observed that the by *codon* partition model and the by gene and codon (*combined*) partition models performed most similar. This corroborates the finding that the codon positions show more heterogeneity in the substitution process than among genes and accounting for this heterogeneity in the substitution process is crucial for robust phylogenetic inference.

**Figure 4.**
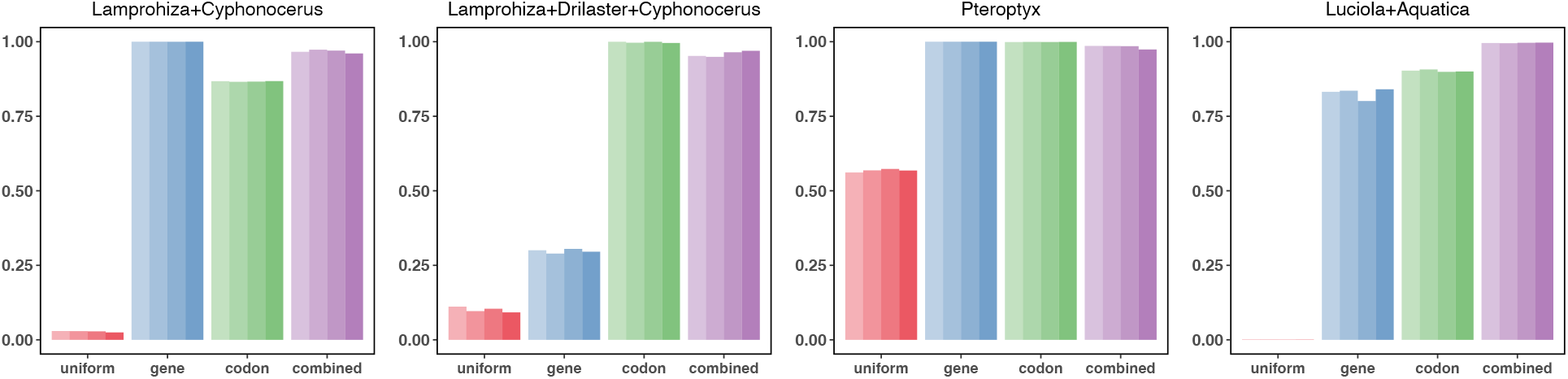
Posterior probabilities of selected clades being monophyletic for different partition models. We performed phylogenetic inference on concatenated alignments assuming four different partition models: uniform, by gene, by codon position, and by codon position and gene (combined). Each analyses was run four times to ensure MCMC convergence. We then computed for each analysis this posterior probability of pre-defined clades to be monophyletic, showing the strength in support or lack thereof. For all tested clades see Fig. S22.

We additionally observed substantial variation in support for genera or other higher clades between single genes (Figs. S15–S16). The lack of support is often correlated with the length of the genes (e.g., support for *Photinus* being monophyletic) with shorter genes lacking support, for example ATP8 contained only 153 sites, ND3 contained only 351 sites, and ND4L contained only 288 sites. The concatenated alignment analyses show overall better supported phylogenies, both in terms of higher posterior probabilities and recovering of established genera, indicating that single genes might not be informative and powerful enough for phylogenetic inference at this timescale (e.g., the genus *Pteroptyx*, Fig. 4 and S15 & S16). As mitochondrial genomes do not recombine, genes from mitochondrial genomes must follow the same evolutionary history and therefore all gene tree discordance is attributed to gene tree estimation error (Richards et al., 2018). Our results confirm that single gene tree estimates can be imprecise likely to due to lack of information and conclusions based on single gene trees should be considered carefully (Höhna et al., 2025).

### 3.3 Time-calibrated firefly phylogeny and impact of partition model on divergence times

Our time-calibrated phylogenetic inferences show very similar estimates across all four partition model settings (Fig. 5). While we observed substantial difference in substitution rates (Fig. 3 left), we do not observe significant differences in clade ages (Fig. 5). The difference in substitution rates is absorbed by difference in average clock rates (Fig. 3 right). Thus, the divergence times estimates are largely dominated by our fossilized-birth-death process prior that calibrates the phylogeny based on fossil occurrence (i.e., induces a minimum constraint on the clade the fossil is placed in).

**Figure 5.**
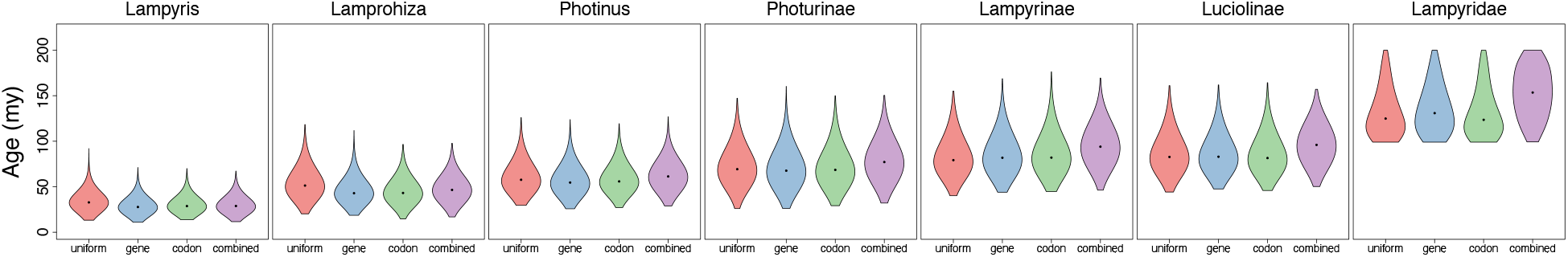
Posterior distributions of clade ages under different partition models. Clade ages are estimated in a concatenated alignment time-calibrated phylogenetic analysis under four different partition strategies: uniform, by gene, by codon position, and combined (both by codon position and gene). Time calibrations were induced through fossil constraints within a fossilized-birth-death process (see Table 1).

At a more nuanced inspection, the *uniform* partition model shows some deviations in clade age estimation for younger clades, while the *combined* (by gene and codon position) partition model shows deviations in clade age estimation for older clades from the other models (Fig. 5). Specifically, the root node of all fireflies is estimated to be older in the *combined* partition model analysis. However, the posterior distribution seems to be most impacted by the prior distribution under this more complex model.

The inferred time-calibrated phylogenies between the different partition models varies in topology in a few ways. First, the topology estimate itself is influenced by the partition model as seen in our unrooted species tree analyses (Fig. 6 and S22–S24). Second, the *uniform* partition model inferred a different position of the root. In the *uniform* partitioned analyses, *Stenocladius* is the single outgroup genus to all other fireflies. This rooting splits the subfamily Ototretinae as paraphyletic. For the partitioned data analyses, we obtained Photurinae and Lampyrinae as one subtree and all other fireflies (e.g., Luciolinae, Lamprohizinzae, Cyphonocerniae and Ototretinae) as the second subtree (Fig. 6).

**Figure 6.**
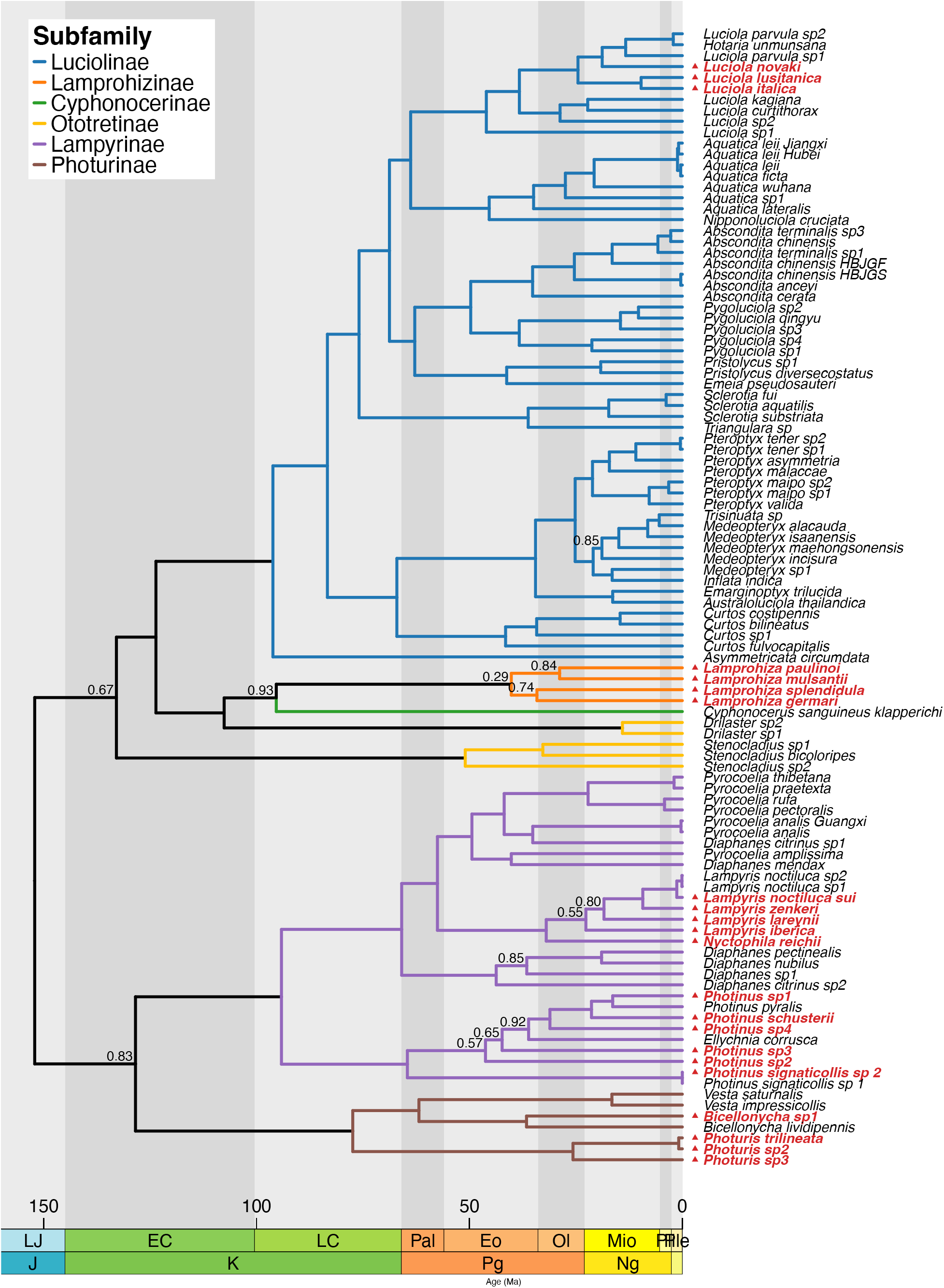
Time-calibrated phylogeny of Lampyridae. Highlighted in red are the 22 newly sequenced firefly species. We colored the phylogeny by the sub-family (see legend). Nodes with a pos2t3erior probability of PP < 0.95 are indicates by their respective posterior probabilities. The phylogeny was plotted using the `R` package `RevGadgets` (Tribble et al., 2022). Here we show the results from the combined partition model analysis.

## 4 Discussion

In this study we sequenced 22 new mitochondrial genomes of 22 different firefly species from Europe (France, Germany, Greece, Italy, Spain and Switzerland) and Nuclear Central America (Guatemala). We explored the impact of partitioning the mitochondrial genome using four strategies: *uniform*, by *gene*, by *codon* position, and *combined* by both gene and codon position. More specifically, we focused on the impact of these four partition strategies on divergence-time estimation. Furthermore, we explored several key-features of phylogenetic inference from mitochondrial genomes, such as the informativeness of single loci compared with concatenation of all loci. We summarize and discuss several of our key findings here.

### 4.1 The impact of partition models on substitution rates and divergence times

We observed the strongest difference in substitution rates among different types of genes (rRNA versus protein coding genes) and among codon positions (Fig. 2). On the one hand, it might be tempting to recommend using a partition model with four data subsets: by gene type and codon position, as done in our by *codon* model. Such a partition model has the clear advantage to be computationally manageable as the number of partitions does not necessarily grow when more loci are added (Shapiro et al., 2006). However, we observed that there is some interaction between the gene and codon position (Fig. 2), which leads to different substitution rate and clock rate estimates (Fig. 3). Nevertheless, this *combined* partition model can become computationally demanding. Therefore, we recommend to partition the data at least by gene type and codon position, and possibly explore some more fine-grained partition models.

While the partition models had a substantial impact on the estimated substitution rates and tree length, these models had only a minor impact on divergence time estimation. Disregarding the impact on the inferred tree topology, it seems to matter little for divergence time estimation which partition model to chose (see also (Angelis et al., 2018)). The difference in substitution rates is completely absorbed by the clock rates (Fig. 3 and 5) and estimates of divergence times are primarily driven by the time-calibration strategy. On the one hand, our observations can be interpreted such that the partition model does not matter if we have reliably estimated the tree topology. On the other hand, our results highlight the interactions of parameter-rich models (highly partition models with among partition rate variation together with relaxed clock models) where naturally the informativeness of the data is reduced.

Estimates of clock rates are affected by the choice of partition model due to their interaction (i.e., higher substitution rates, as evidenced by larger tree lengths, resulted in higher clock rates because divergence time estimates remain roughly equivalent). Our clock rate estimates are in general slower than previously reported for mitochondrial genomes in coleoptera (Pons et al., 2010) and the *combined* partition model comes closest to the previously published clock rate (Fig. 3). This insight can play a major role when clock rates are used to time-calibrate phylogenies when no other calibrations are available.

In our study we observed that tree lengths of single gene phylogenetic analyses are conspicuously close to the prior expectation, especially when partitioning by codon position (Fig. 2). This means that for single mitochondrial genes there is not enough information in the data at the timescale of fireflies to overpower the prior (Yang and Rannala, 2005). Contrary to this, multi-locus concatenated analyses show clear deviations from the prior, i.e., the posterior distributions did not contain the prior mean (Fig. 3). It might therefore be questionable how reliable we can infer timescales from single gene trees, or robustly inferring variations in ages of single gene trees within a multi-species coalescent analysis, as these are likely entirely driven by choices of the prior.

### 4.2 Variation of substitution rates across genes and codon positions

Our results showed that there is up to 3-fold difference in substitution rates among different genes, and much higher differences between codon positions (up to 14-fold). Nevertheless, for the same codon position, we observe at most again a 3-fold variation (Fig. 2). When we incorporate among partition rate variation into our phylogenetic models, then we assume *a priori* that any amount of partition rate variation is equally probable due to the flat Dirichlet prior distribution (Dos Reis et al., 2014; Höhna et al., 2017). Thus, our results shed some light into how among partition rate variation is distributed in empirical data, although more studies would be helpful. From a modeling perspective, we might find a better trade-off than either assuming no among partition rate variation (the *uniform* partition strategy) or assuming that all partition rates are equally probable *a priori* (the flat Dirichlet prior). For example, using a lognormal or gamma prior distribution on among partition rate variation with a 3-fold variance might provide a better fit, although care needs to be taken to design such a model appropriately (Dos Reis et al., 2014). In the early days of statistical phylogenetic models, it was suggested that the +Γ for among site rate variation is sufficient and will capture variation among sites within a partition (Yang, 1993, 1994; Rannala and Yang, 2008). However, our results show that this is not the case as otherwise partitioning by codon position should have no impact on the overall substitution or clock rate because we always used the +Γ within in each partition. Each data subset additionally showed among character rate variation (+Γ), thus not all variation in substitution rates is absorbed by among partition rate variation.

### 4.3 Robustness of single gene trees

Our single gene tree estimates showed substantial uncertainty and lack of power even for very commonly applied standard genes such as cytochrome c oxidase subunit 1 gene (CO1), cytochrome b (CYTB), and NADH dehydrogenase 1 protein (Duchêne et al., 2011). The multi-locus concatenated analyses, on the other hand, were more robust, questioning whether single loci are reliable enough for robust phylogenetic inference (Scornavacca and Galtier, 2017). This observation has two main implications. First, given today’s reduction in sequencing cost, we strongly recommend to sequence more loci and, if mitochondrial loci are sequenced, then sequencing the full mitochondrial genome for robust phylogenetic inference (Duchêne et al., 2011). Second, single gene-tree inferences appear unreliable, especially for mitochondrial genes (Richards et al., 2018; Toups et al., 2025), which had also been observed for anchored hybrid enrichment data in fireflies (Höhna et al., 2025). Since the mitochondrial genome does not recombine, all gene tree discordance can be attributed to gene tree estimation error instead of actual biological processes such as deep coalescence (incomplete lineage sorting), gene paralogs due to duplication and losses, and horizontal gene transfer (Maddison and Knowles, 2006; Degnan and Rosenberg, 2009). Our results show that single gene tree estimate can be flawed, especially when the underlying loci are short, and inference about causes of gene tree discordance has to be considered carefully.

### 4.4 Time-calibrated firefly phylogeny

Our time-calibrated phylogeny agrees with the overall classification of subfamilies with the exception of Ototretinae. Among the subfamilies, we recovered Lamprohizinae as sister to Cyphonicerinae and Ototretinae, all on the same side of the phylogeny as Luciolinae. Several previous studies placed Lamprohizinae on the adjacent subtree together with Lampyrinae and Photurinae (Stanger-Hall et al., 2007; Martin et al., 2017, 2019; Höhna et al., 2025). Our conclusions show that the early relationship among fireflies, especially the position of Lamprohizinae, Cyphonicerinae and Ototretinae, needs further investigation. Given the discordance between mitochondrial and multi-locus nuclear phylogenies, the most promising approach seems to be utilizing new full genome sequences (Catalán et al., 2024, 2025a).

We recovered most genera to be monophyletic and well resolved (Fig. 6 and Fig. S21). The genus *Diaphanes* was recovered as paraphyletic and requires careful reconsideration (Chen et al., 2019). Specifically, *Diaphanes citrinus sp1* and *Diaphanes mendax* (Chen et al., 2019) cluster consistently together within *Pyrocoelia*. We could also confidently place *Vesta* within Photurinae as sister to *Bicellonycha* (Chen et al., 2019).

From the 22 sequenced firefly mitochondrial genomes, 18 species had not been placed before in a phylogenetic context. For European fireflies, we added resolution on species relationships and divergence estimates within the genera *Luciola, Lamprohiza* and *Lampyris*. This also applies to the genera *Photinus, Bicellonycha* and *Photuris*, where mostly North American species have been placed in a phylogeny. Our new firefly species from Guatemala depict a considerable phylogenetic diversity (Catalán et al., 2025b). Furthermore, these Guatemalan firefly samples elucidate the origin of North American fireflies, for example, the North American firefly *Photinus pyralis* is nested within our Guatemalan *Photinus* samples. To draw further conclusions, we would need more mitochondria samples from both geographical regions (Catalan et al., 2022).

Our age estimates push back the origin of *Photinus* towards the KPg boundary (Fig. 5 and Fig. 6), whereas previous estimates reported *Photinus* to originate around 40 MYA (Catalan et al., 2022; Powell et al., 2022; Höhna et al., 2025). Our age estimates of *Lamprohiza* and *Lampyris* are also slightly pushed back, probably due to more extensive taxon sampling. These older time estimates of these genera have an impact on some subfamilies (e.g., Photurinae) but not on others (e.g., Lampyrinae). However, our study has a much denser taxon sampling for Luciolinae and much lighter taxon sampling for Lampyrinae compared with the two previous time-calibrated phylogenies (Powell et al., 2022; Höhna et al., 2025).

There are two main conclusions from our new divergence time estimation. First, a more complete taxon sampling is needed to obtain a more precise picture of the timing of divergence in fireflies. Second, better and more precise fossil calibrations are need to refine our understanding of the timing of firefly diversification, especially as our current estimates have comparably large credible intervals (Fig. 5). The most promising approach could be to obtain more confident calibrations about the root age of fireflies which would have the strongest impact on divergence times of subfamilies and genera.

## 5 Conclusion

In this study we explored the impact of partition models on time-calibrated phylogenetic inference from mitochondrial genomes. In our case study, we used a firefly dataset of 22 new sequenced mitochondrial genomes and 82 published mitochondrial genomes. We show importance to partition the data at least by gene type and codon position as these exhibit significantly heterogeneous substitution processes impacting phylogenetic inference. Although divergence times are almost not affected by the choice of partition model, the underlying topology including rooting of the phylogeny can be affected as well as the clock rate, complicating comparison and clock-rate calibrations among studies. Finally, we show that among partition rate variation between mitochondrial genes is at most 3-fold, and the strongest variation is among codon positions. These insights can lead to development of more realistic models of among partition rate variation compared with the rather simplistic flat Dirichlet prior that is exclusively used today.

## Acknowledgements

This study was supported by a Deutsche Forschungsgemeinschaft priority program “The genomic basis of evolutionary innovations” (SPP2349; Project No. 503272152 awarded to SH [HO 6201/3-1] and AC [CA 2207/4-1]).

