## Supplementary Material for "Impact of Partition Models on Phylogenetic Inference and Divergence Times of Lampyridae from Mitochondrial Genomes"

### Supporting Information

[1,2]Sebastian Höhna [1,2]Haoqing Du [3]Ana Catalán
[1]*GeoBio-Center LMU, Ludwig-Maximilians-Universität München, 80333 Munich, Germany* [2]*Department of Earth and Environmental Sciences, Paleontology & Geobiology, Ludwig-Maximilians-Universität München, 80333 Munich, Germany*
[3]*Division of Evolutionary Biology, Ludwig-Maximilians-Universität München*

### S1 Lampyridae Mitochondrial Genomes

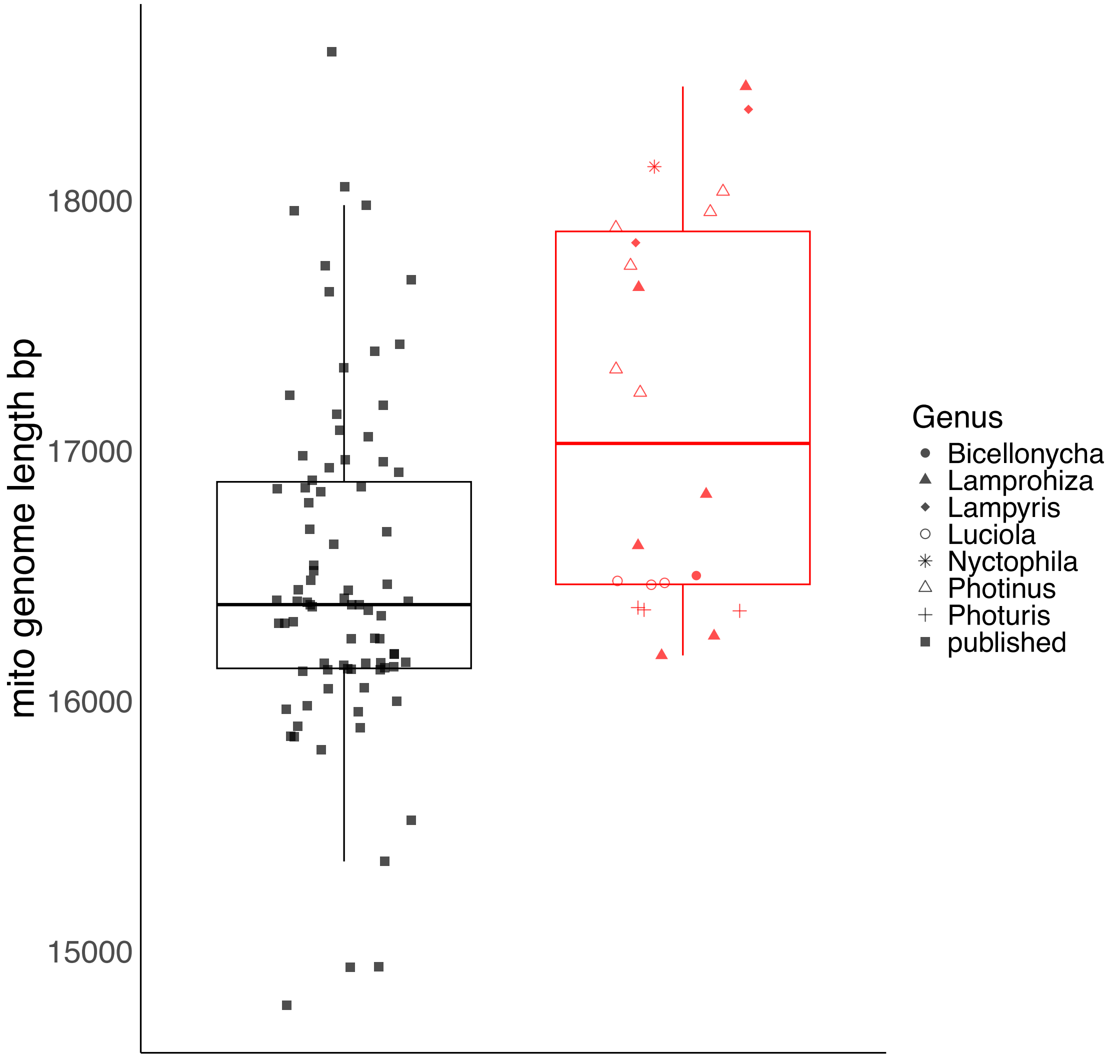

**Figure S1: Distribution of mitochondrial genome lengths.** Left (black): lengths of published mitochondrial genomes. Right (red): lengths of newly sequenced mitochondrial genomes. We additionally coded the new sequences by genus (see legend).

**Table S1: Genera, Tribes and Sub-families used to assess gene-tree error and clade ages.**

| Species | Tribe | Subfamily | Accession | Reference |
|---|---|---|---|---|
| Abscondita anceyi sp1 | - | Luciolinae | NC_039706 | Hu and Fu (2018a) |
| Abscondita anceyi sp2 | - | Luciolinae | MT554383 | |
| Abscondita cerata | - | Luciolinae | MW751423 | Wang et al. (2021) |
| Abscondita chinensis HBJGF | - | Luciolinae | OL998464 | |
| Abscondita chinensis HBJGS | - | Luciolinae | NC_068007 | |
| Abscondita chinensis | - | Luciolinae | OM802863 | |
| Abscondita terminalis sp1 | - | Luciolinae | NC_044776 | Chen et al. (2019) |
| Abscondita terminalis sp2 | - | Luciolinae | MN722653 | Liu and Fu (2020) |
| Abscondita terminalis sp3 | - | Luciolinae | OM802860 | |
| Aquatica ficta | - | Luciolinae | NC_035060 | Wang et al. (2017) |
| Aquatica lateralis | - | Luciolinae | OM135506 | |
| Aquatica leii | - | Luciolinae | NC_025276 | |
| Aquatica leii Jiangxi | - | Luciolinae | OM743434 | |
| Aquatica leii Hubei | - | Luciolinae | OM743435 | |
| Aquatica sp1 | - | Luciolinae | OM135505 | |
| Aquatica wuhana | - | Luciolinae | NC_035061 | Wang et al. (2017) |
| Asymmetricata circumdata | - | Luciolinae | NC_032062 | |
| Australoluciola thailandica | - | Luciolinae | NC_084295 | |
| Bicellonycha lividipennis | - | Photurinae | NC_030060 | Amaral et al. (2016) |
| Bicellonycha sp1 | - | Photurinae | | this study |
| Curtos bilineatus | - | Luciolinae | NC_044789 | Chen et al. (2019) |
| Curtos costipennis | - | Luciolinae | MK609965 | Zhang and Fu (2019) |
| Curtos fulvocapitalis | - | Luciolinae | NC_058281 | |
| Curtos sp1 | - | Luciolinae | OP747313 | |
| Cyphonocerus sanguineus klapperichi | - | Psilocladinae | MW365445 | |
| Diaphanes citrinus sp1 | Lampyrini | Lampyrinae | MK292103 | Chen et al. (2019) |
| Diaphanes citrinus sp2 | Lampyrini | Lampyrinae | NC_051869 | Yang and Fu (2019) |
| Diaphanes mendax | Lampyrini | Lampyrinae | NC_044791 | Chen et al. (2019) |
| Diaphanes nubilus | Lampyrini | Lampyrinae | NC_044787 | Chen et al. (2019) |
| Diaphanes pectinealis | Lampyrini | Lampyrinae | NC_044793 | Chen et al. (2019) |
| Diaphanes sp1 | Lampyrini | Lampyrinae | MK292095 | Chen et al. (2019) |
| Drilaster sp1 | - | Ototretinae | MZ457901 | |
| Drilaster sp2 | - | Ototretinae | MK292100 | Chen et al. (2019) |
| Ellychnia corrusca | Photinini | Lampyrinae | MG242622 | |
| Emarginoptyx trilucida | - | Luciolinae | NC_067971 | |
| Emeia pseudosauteri | - | Luciolinae | MN722654 | Liu and Fu (2020) |
| Inflata indica | - | Luciolinae | NC_039700 | Sriboonlert and Wonnapinij (2019) |
| Lamprigera yunnana | Photinini | Lampyrinae | NC_044786 | Chen et al. (2019) |
| Lamprohiza germari | - | Lamprohizinae | | this study |
| Lamprohiza mulsantii | - | Lamprohizinae | | this study |
| Lamprohiza paulinoi | - | Lamprohizinae | | this study |
| Lamprohiza splendidula | - | Lamprohizinae | | this study |
| Lampyris iberica | Lampyrini | Lampyrinae | | this study |
| Lampyris lareynii | Lampyrini | Lampyrinae | | this study |
| Lampyris noctiluca sp1 | Lampyrini | Lampyrinae | KX087302 | |
| Lampyris noctiluca sp2 | Lampyrini | Lampyrinae | MN122858 | |
| Lampyris noctiluca sui | Lampyrini | Lampyrinae | | this study |
| Lampyris zenkeri | Lampyrini | Lampyrinae | | this study |
| Luciola curtithorax | - | Luciolinae | NC_038225 | Hu and Fu (2018b) |
| Luciola italica | - | Luciolinae | | this study |
| Luciola kagiana | - | Luciolinae | NC_072664 | |
| Luciola lusitanica | - | Luciolinae | | this study |
| Luciola novaki | - | Luciolinae | | this study |
| Luciola parvula sp1 | - | Luciolinae | LC677171 | |
| Luciola parvula sp2 | - | Luciolinae | NC_067969 | |
| Luciola sp1 | - | Luciolinae | OP747314 | |
| Luciola sp2 | - | Luciolinae | OP747315 | |
| Luciola substriata | - | Luciolinae | NC_027176 | Mu et al. (2016) |
| Hotaria unmunsana | - | Luciolinae | NC_050947 | Kim et al. (2020) |
| Medeopteryx alacauda | - | Luciolinae | NC_084296 | |
| Medeopteryx incisura | - | Luciolinae | NC_084297 | |
| Medeopteryx isaanensis | - | Luciolinae | NC_084298 | |
| Medeopteryx maehongsonensis | - | Luciolinae | NC_084299 | |
| Medeopteryx sp1 | - | Luciolinae | OL891941 | |
| Nipponoluciola cruciata | - | Luciolinae | OM718717 | |
| Nyctophila reichii | Lampyrini | Lampyrinae | | this study |
| Photinus pyralis | Photinini | Lampyrinae | KY778696 | Fallon et al. (2018) |
| Photinus schusterii | Photinini | Lampyrinae | | this study |
| Photinus signaticollis | Photinini | Lampyrinae | NC_071760 | Koken and Gastineau (2024) |
| Photinus signaticollis sp2 | Photinini | Lampyrinae | | this study |
| Photinus sp1 | Photinini | Lampyrinae | | this study |
| Photinus sp2 | Photinini | Lampyrinae | | this study |
| Photinus sp3 | Photinini | Lampyrinae | | this study |
| Photinus sp4 | Photinini | Lampyrinae | | this study |
| Photuris sp1 | - | Photurinae | | this study |
| Photuris sp2 | - | Photurinae | | this study |
| Photuris sp3 | - | Photurinae | | this study |
| Pristolycus diversecostatus | - | Luciolinae | NC_067973 | |
| Pristolycus sp1 | - | Luciolinae | MK292099 | Chen et al. (2019) |
| Pteroptyx asymmetria | - | Luciolinae | NC_084300 | |
| Pteroptyx maipo sp1 | - | Luciolinae | NC_036353 | Fan and Fu (2017) |
| Pteroptyx maipo sp2 | - | Luciolinae | OM802859 | |
| Pteroptyx malaccae | - | Luciolinae | NC_084301 | |
| Pteroptyx tener sp1 | - | Luciolinae | NC_067972 | |
| Pteroptyx tener sp2 | - | Luciolinae | OP747322 | |
| Pteroptyx valida | - | Luciolinae | NC_084302 | |
| Pygoluciola qingyu | - | Luciolinae | NC_057261 | Liu and Fu (2020) |
| Pygoluciola sp1 | - | Luciolinae | OM201323 | |
| Pygoluciola sp2 | - | Luciolinae | OM201324 | |
| Pygoluciola sp3 | - | Luciolinae | MZ571356 | |
| Pygoluciola sp4 | - | Luciolinae | OP747324 | |
| Pyrocoelia amplissima | Lampyrini | Lampyrinae | PP935118 | |
| Pyrocoelia analis | Lampyrini | Lampyrinae | OK323960 | Ji and Xu (2022) |
| Pyrocoelia analis Guangxi | Lampyrini | Lampyrinae | NC_068744 | |
| Pyrocoelia pectoralis | Lampyrini | Lampyrinae | NC_085282 | |

### S2 Gene trees

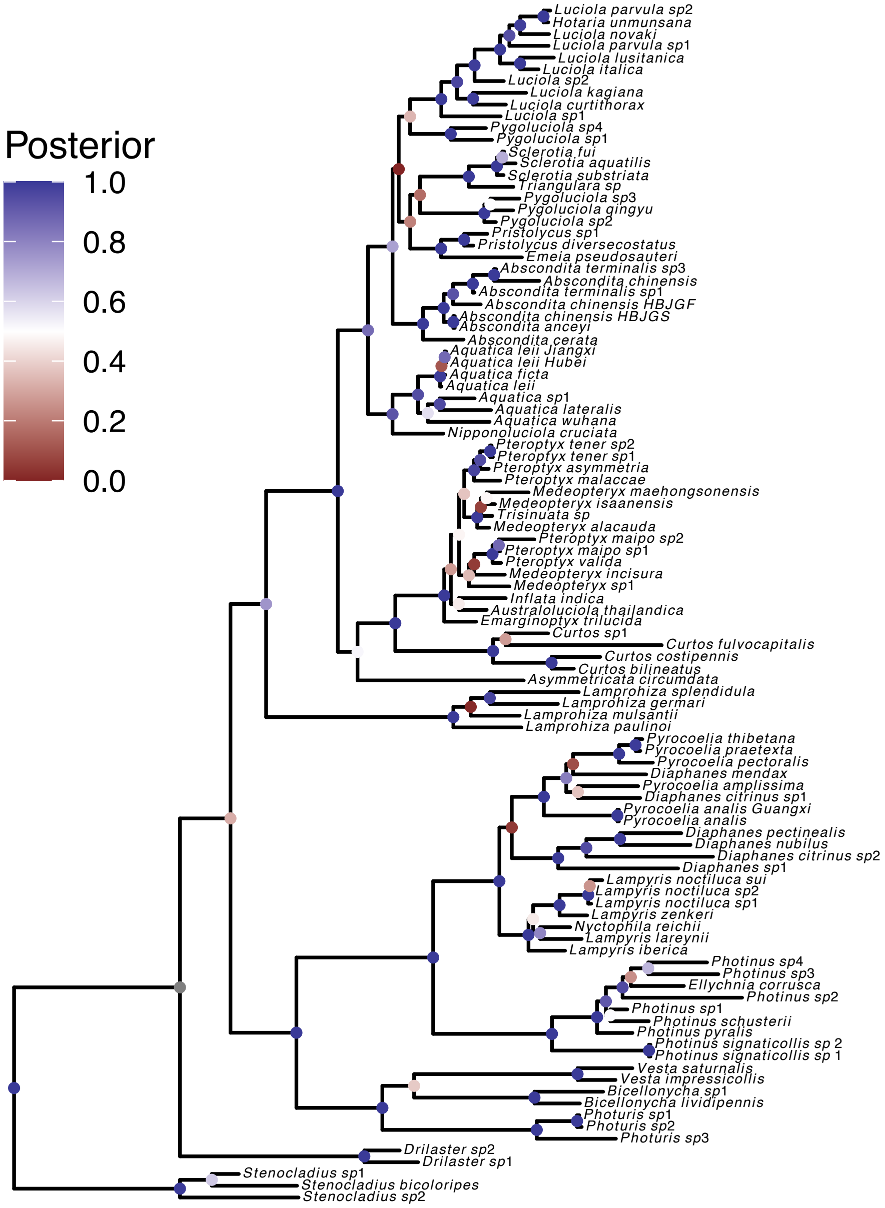

**Figure S2: Maximum a posteriori gene tree for the rRNA 12S locus.** Here we show the *maximum a posteriori* (MAP) gene tree, i.e., the gene tree with the highest posterior probability. Circles at internal nodes are colored based on posterior probabilities. The tree was rooted with *Stenocladius* as outgroup.

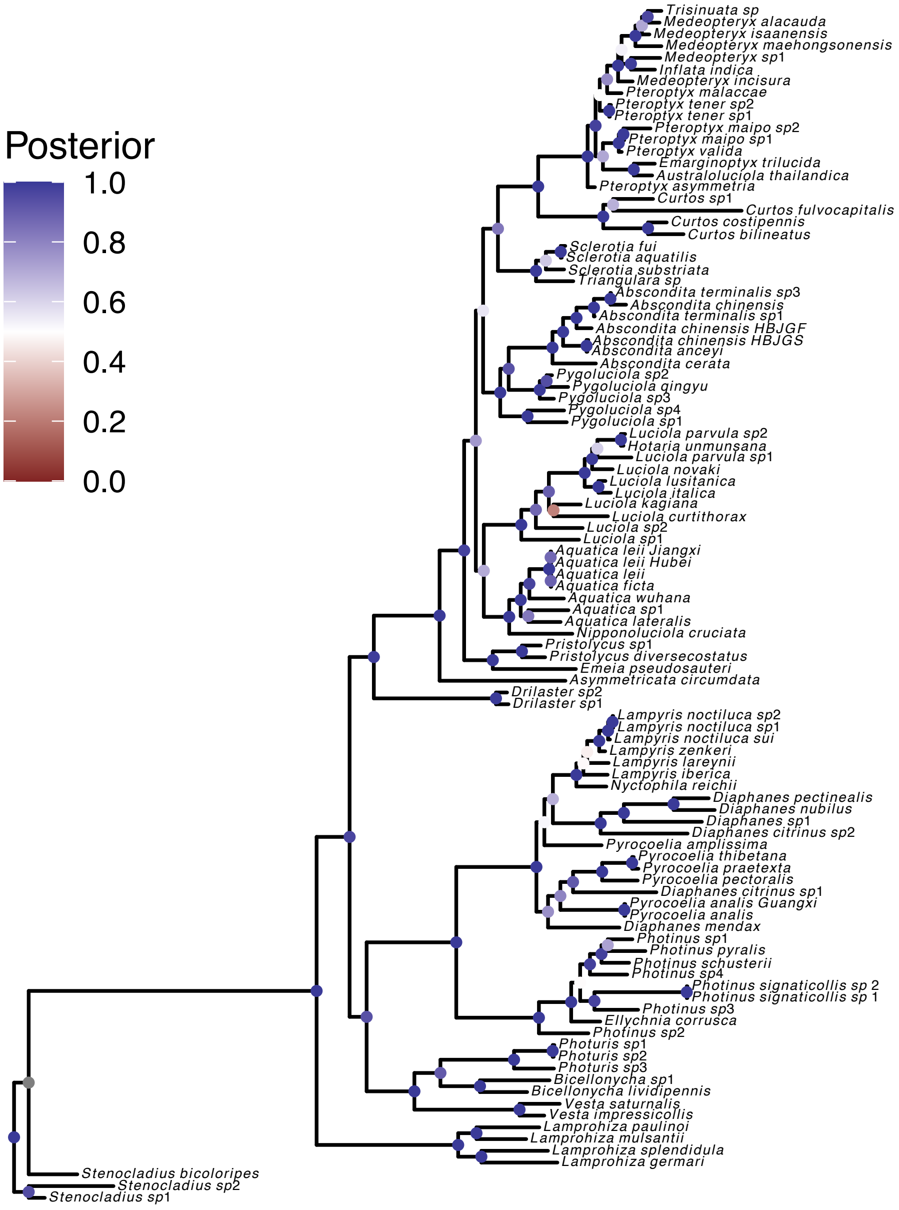

**Figure S3: Maximum a posteriori gene tree for the rRNA 16S locus.** Here we show the *maximum a posteriori* (MAP) gene tree, i.e., the gene tree with the highest posterior probability. Circles at internal nodes are colored based on posterior probabilities. The tree was rooted with *Stenocladius* as outgroup.

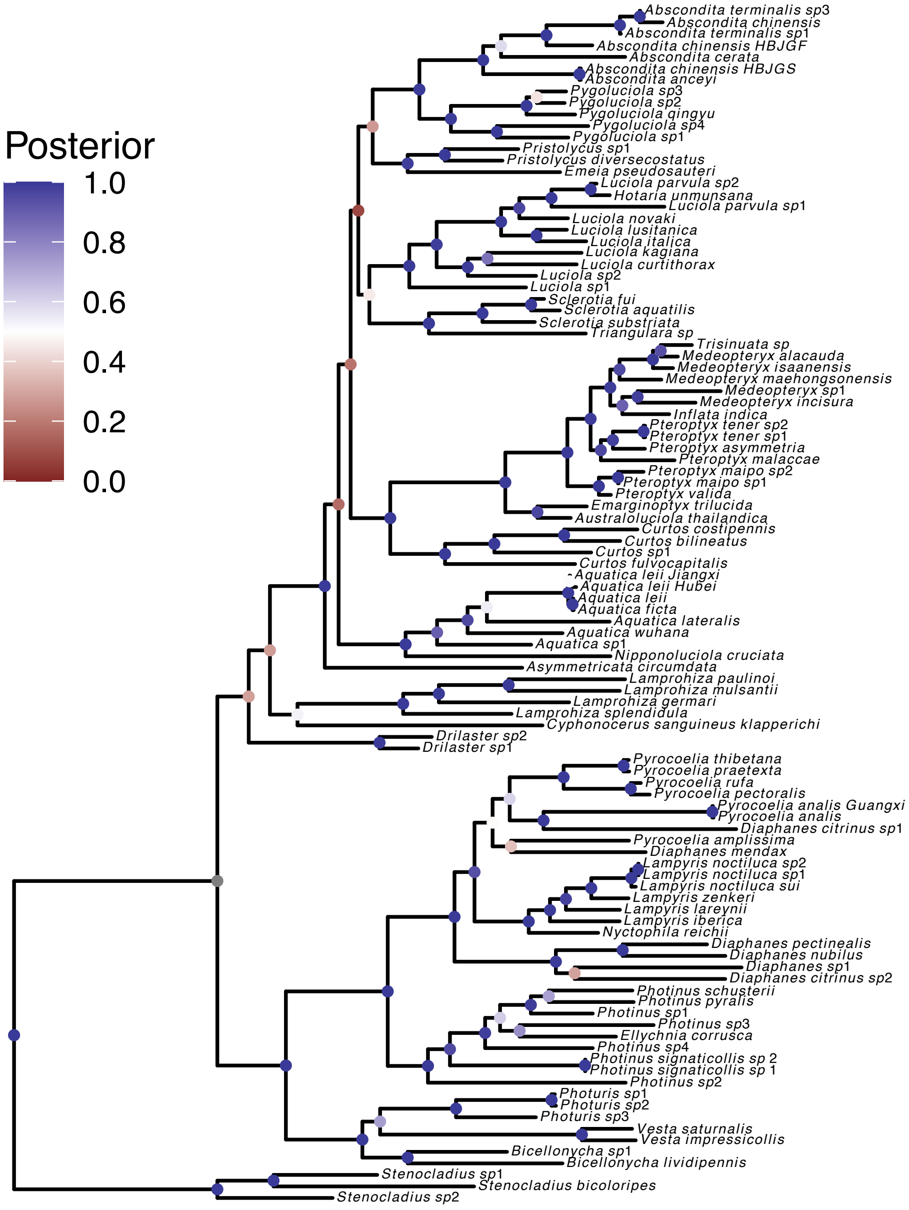

**Figure S4: Maximum a posteriori gene tree for the protein coding gene COX-1 locus.** Here we show the *maximum a posteriori* (MAP) gene tree, i.e., the gene tree with the highest posterior probability. Circles at internal nodes are colored based on posterior probabilities. The tree was rooted with *Stenocladius* as outgroup.

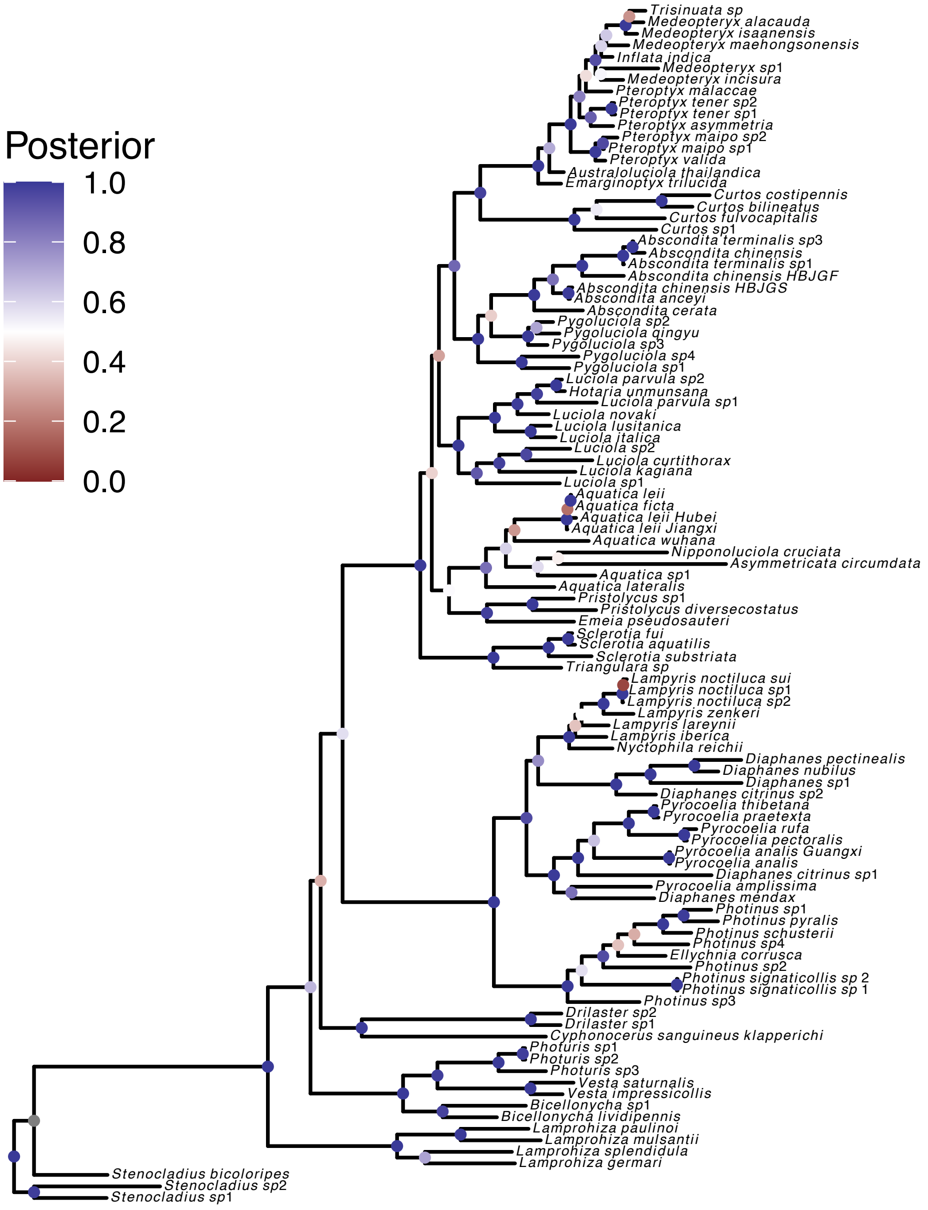

**Figure S5: Maximum a posteriori gene tree for the protein coding gene COX-2 locus.** Here we show the *maximum a posteriori* (MAP) gene tree, i.e., the gene tree with the highest posterior probability. Circles at internal nodes are colored based on posterior probabilities. The tree was rooted with *Stenocladius* as outgroup.

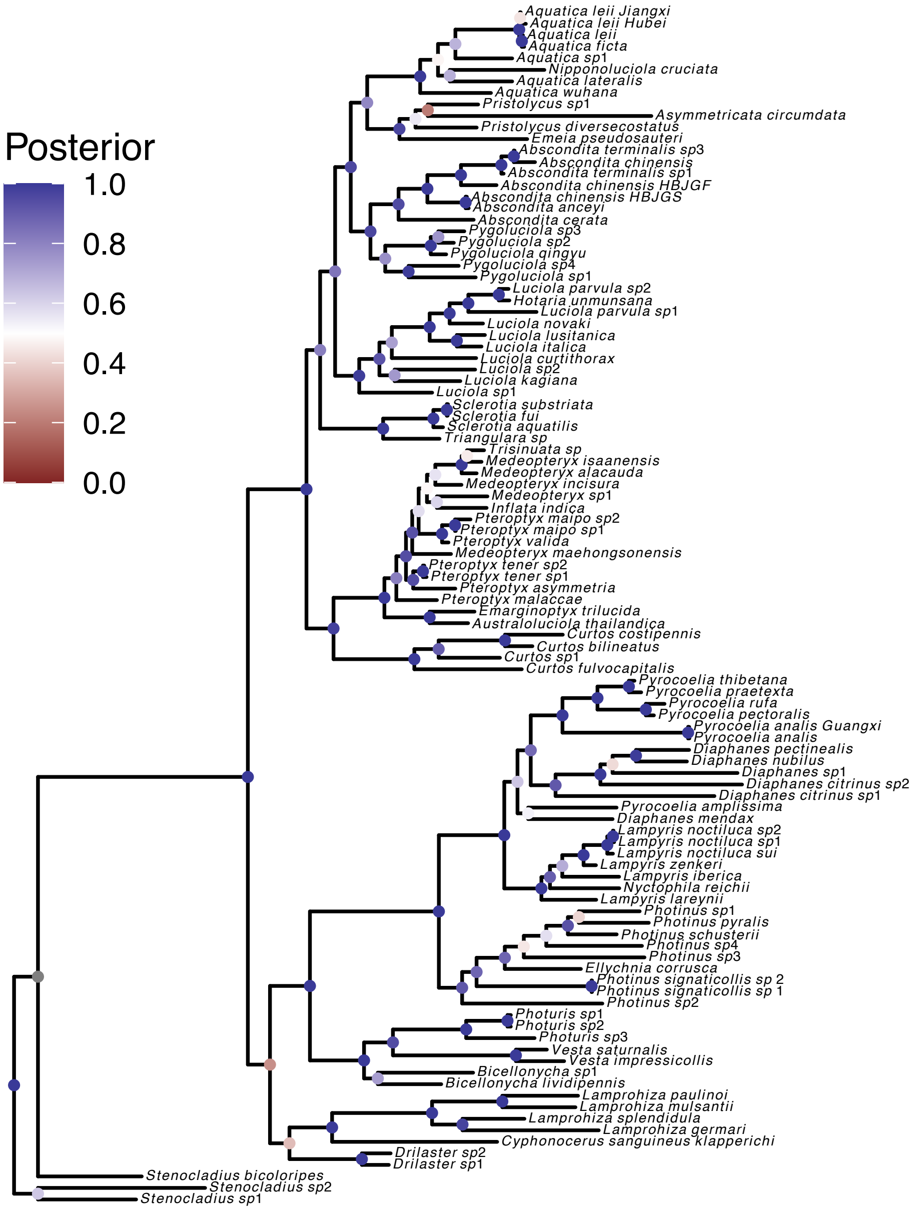

**Figure S6: Maximum a posteriori gene tree for the protein coding gene COX-3 locus.** Here we show the *maximum a posteriori* (MAP) gene tree, i.e., the gene tree with the highest posterior probability. Circles at internal nodes are colored based on posterior probabilities. The tree was rooted with *Stenocladius* as outgroup.

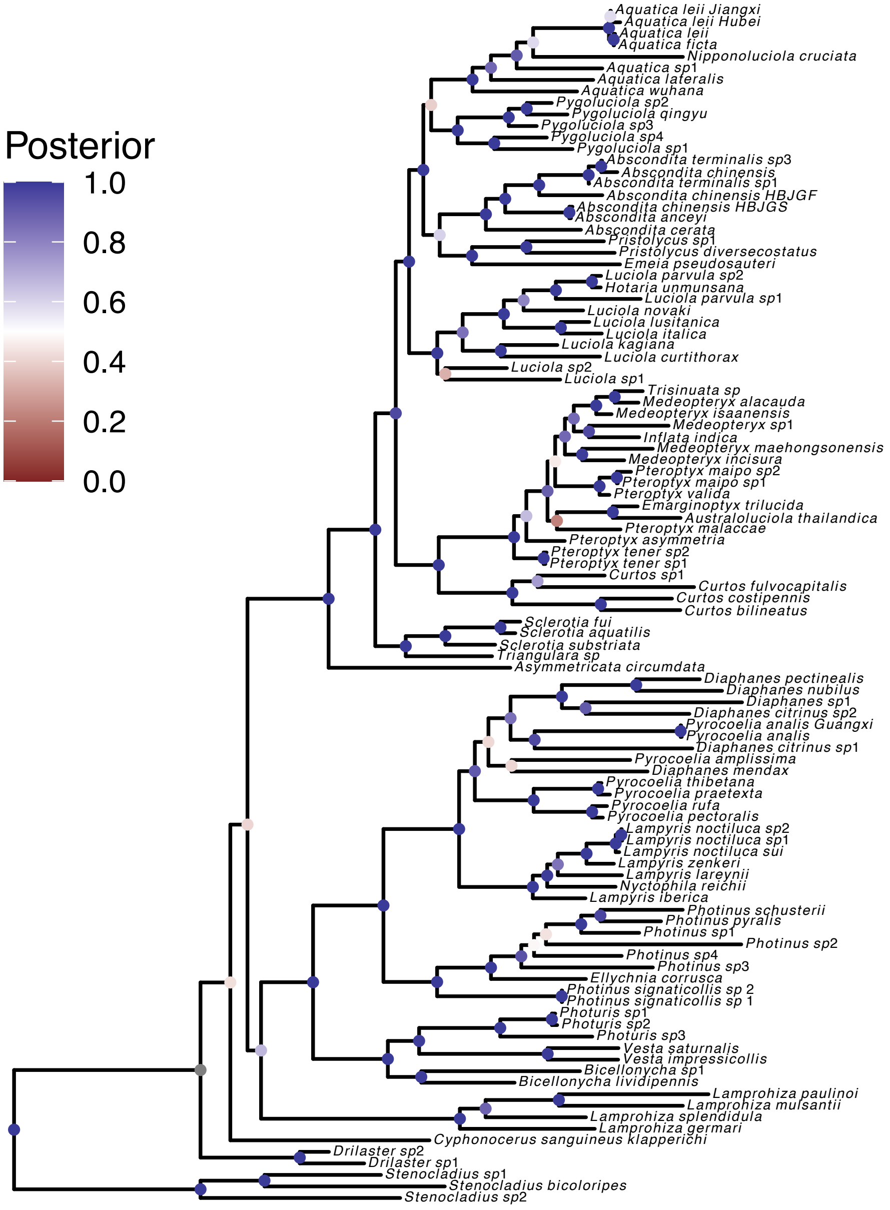

**Figure S7: Maximum a posteriori gene tree for the protein coding gene cytochrome b locus.** Here we show the *maximum a posteriori* (MAP) gene tree, i.e., the gene tree with the highest posterior probability. Circles at internal nodes are colored based on posterior probabilities. The tree was rooted with *Stenocladius* as outgroup.

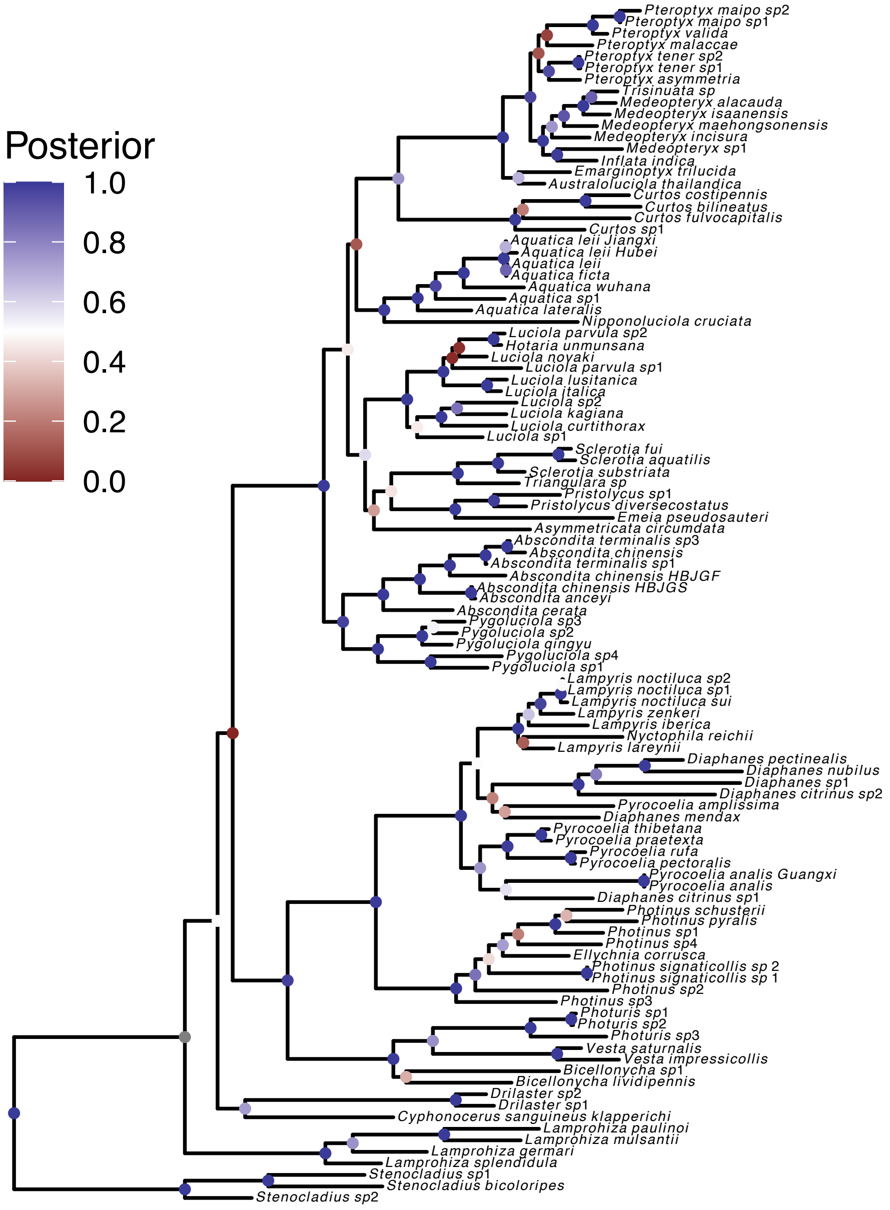

**Figure S8: Maximum a posteriori gene tree for the protein coding gene NADH-1 locus.** Here we show the *maximum a posteriori* (MAP) gene tree, i.e., the gene tree with the highest posterior probability. Circles at internal nodes are colored based on posterior probabilities. The tree was rooted with *Stenocladius* as outgroup.

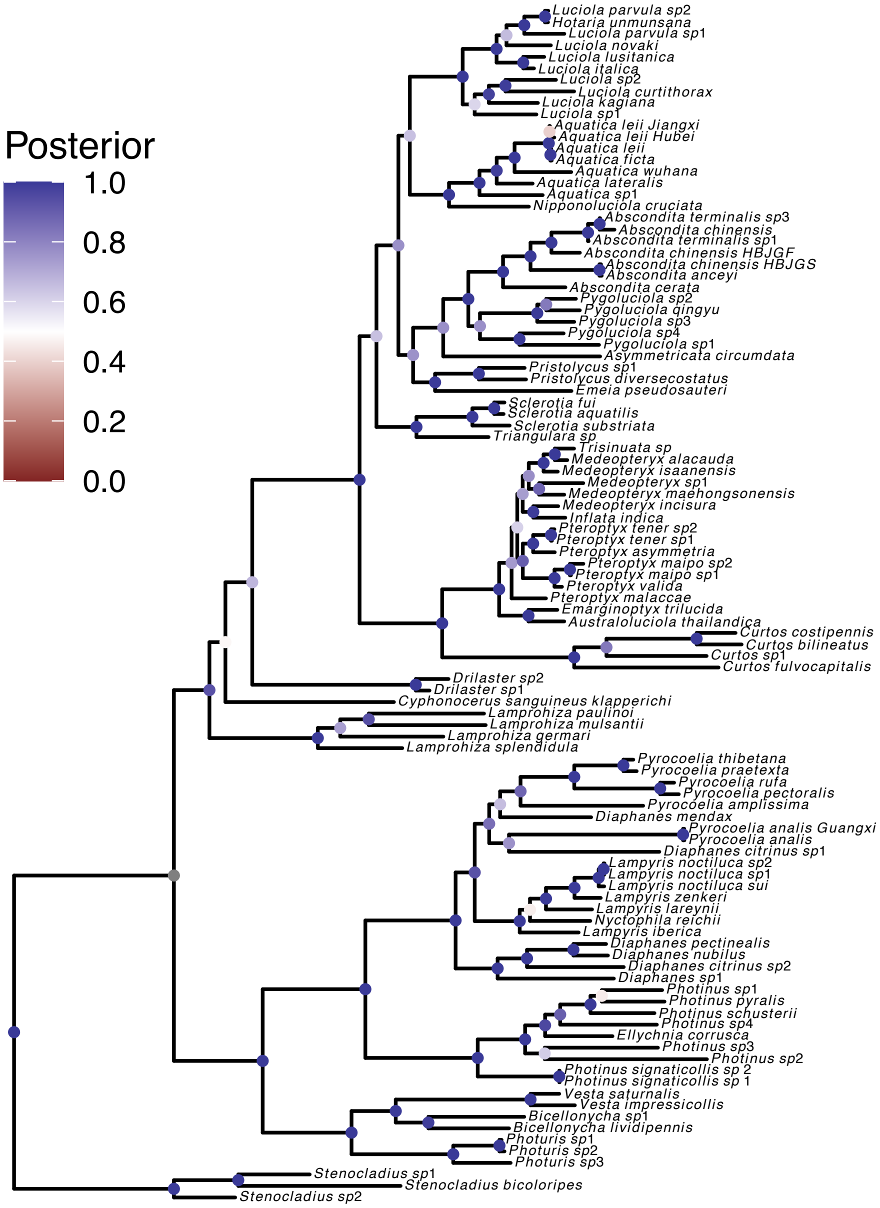

**Figure S9: Maximum a posteriori gene tree for the protein coding gene NADH-2 locus.** Here we show the *maximum a posteriori* (MAP) gene tree, i.e., the gene tree with the highest posterior probability. Circles at internal nodes are colored based on posterior probabilities. The tree was rooted with *Stenocladius* as outgroup.

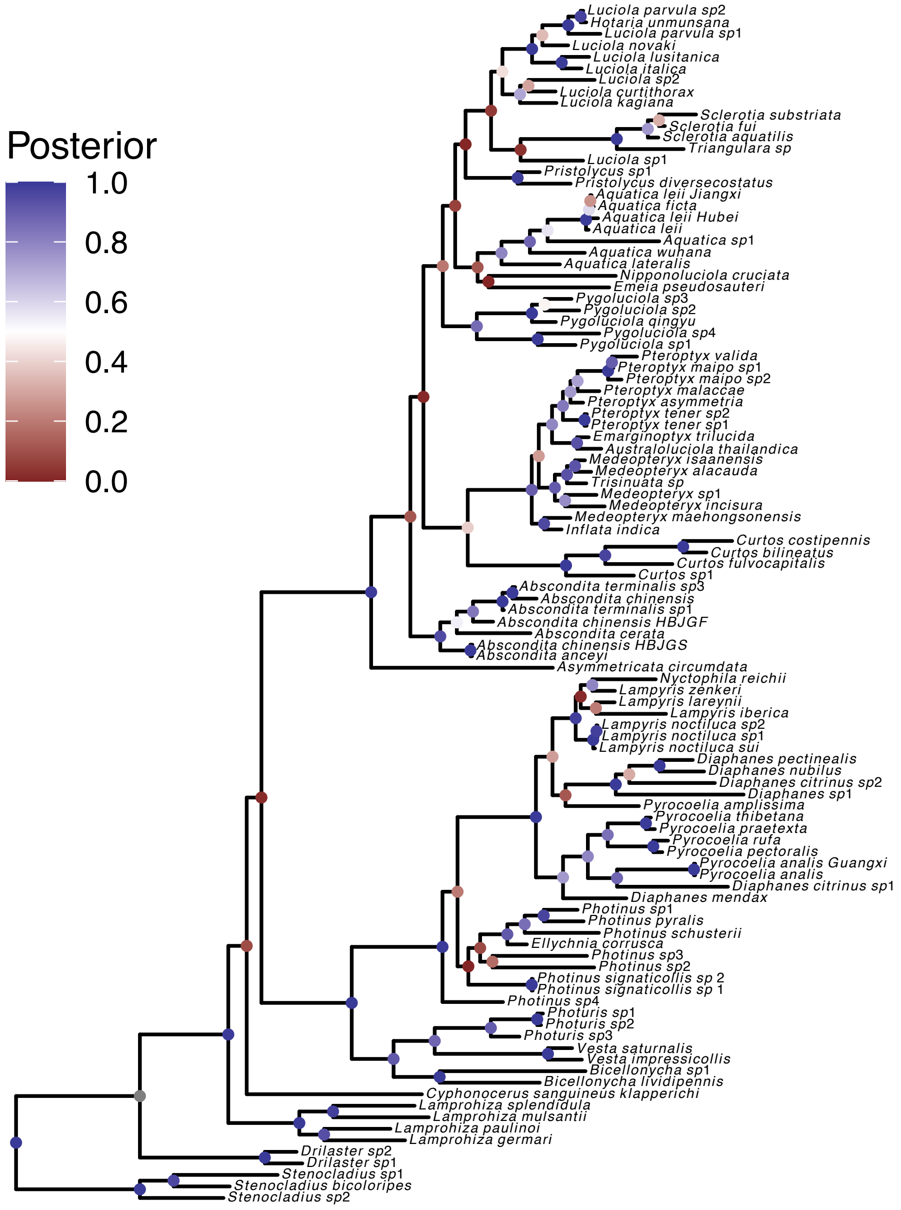

**Figure S10: Maximum a posteriori gene tree for the protein coding gene NADH-3 locus.** Here we show the *maximum a posteriori* (MAP) gene tree, i.e., the gene tree with the highest posterior probability. Circles at internal nodes are colored based on posterior probabilities. The tree was rooted with *Stenocladius* as outgroup.

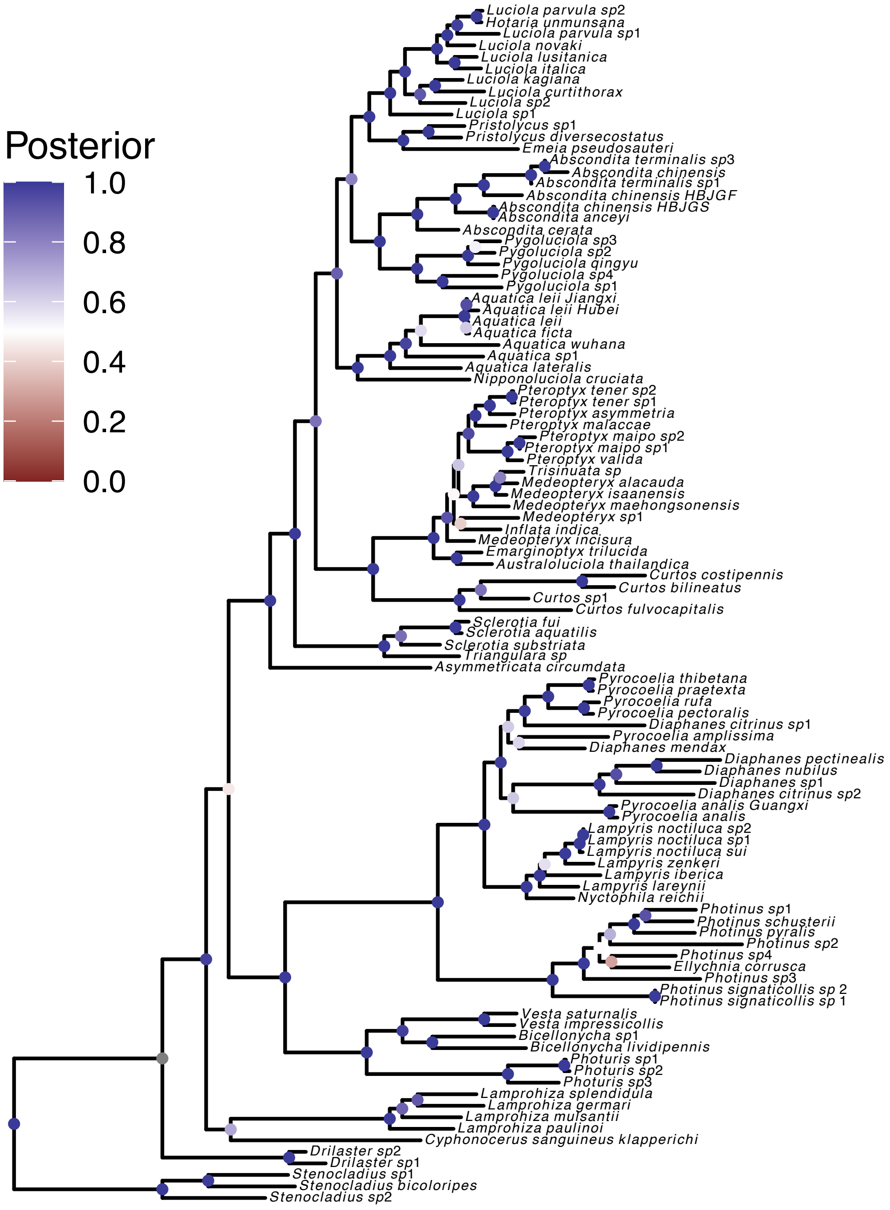

**Figure S11: Maximum a posteriori gene tree for the protein coding gene NADH-4 locus.** Here we show the *maximum a posteriori* (MAP) gene tree, i.e., the gene tree with the highest posterior probability. Circles at internal nodes are colored based on posterior probabilities. The tree was rooted with *Stenocladius* as outgroup.

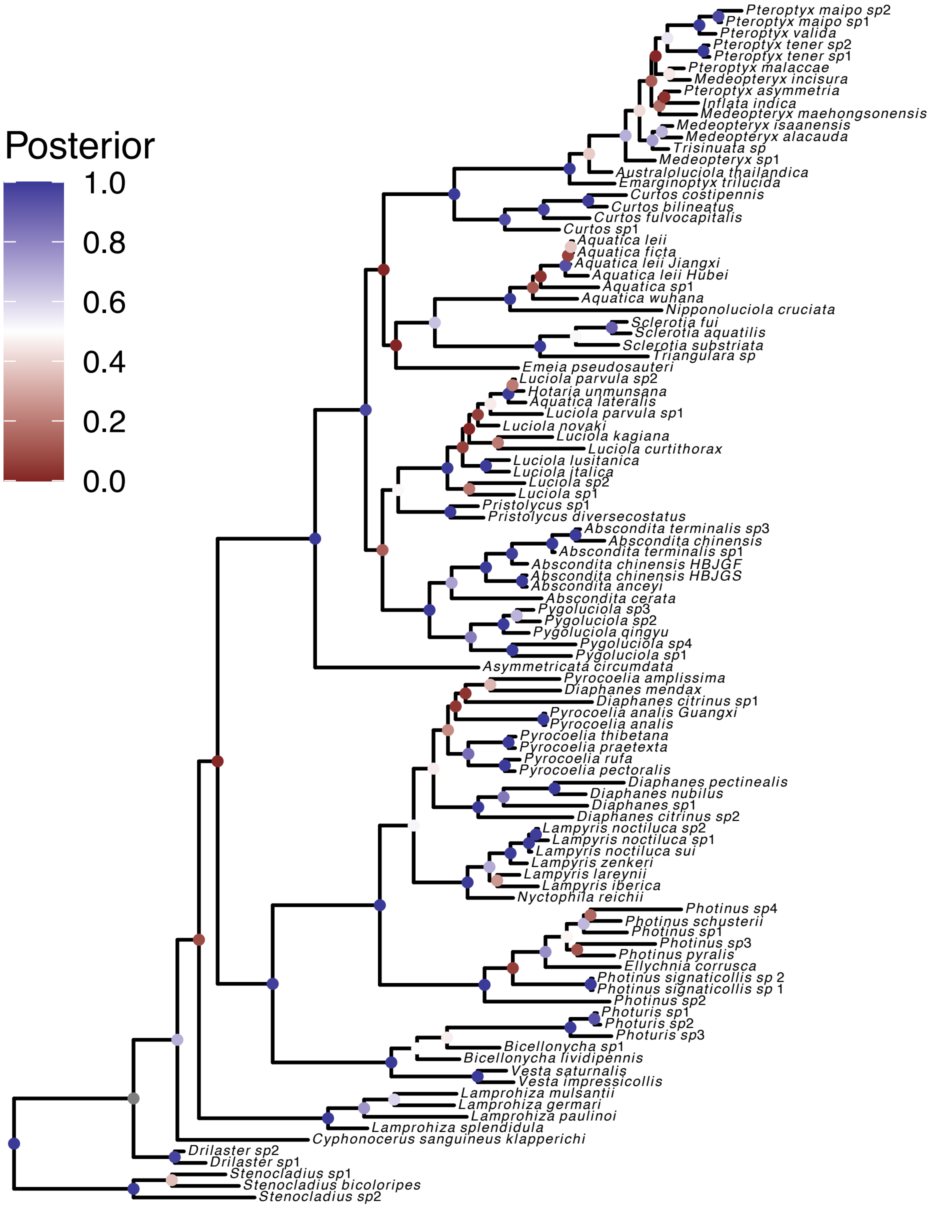

**Figure S12: Maximum a posteriori gene tree for the protein coding gene NADH-4L locus.** Here we show the *maximum a posteriori* (MAP) gene tree, i.e., the gene tree with the highest posterior probability. Circles at internal nodes are colored based on posterior probabilities. The tree was rooted with *Stenocladius* as outgroup.

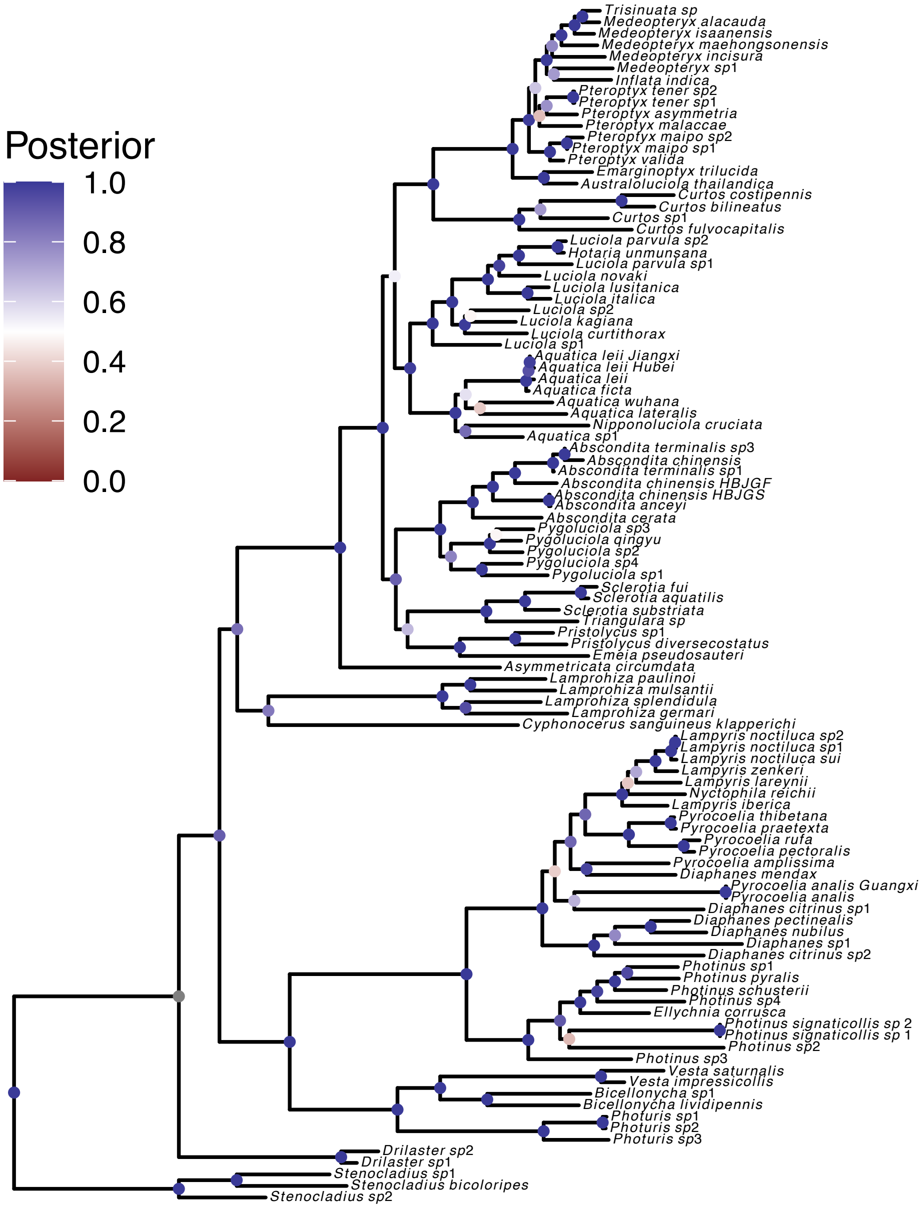

**Figure S13: Maximum a posteriori gene tree for the protein coding gene NADH-5 locus.** Here we show the *maximum a posteriori* (MAP) gene tree, i.e., the gene tree with the highest posterior probability. Circles at internal nodes are colored based on posterior probabilities. The tree was rooted with *Stenocladius* as outgroup.

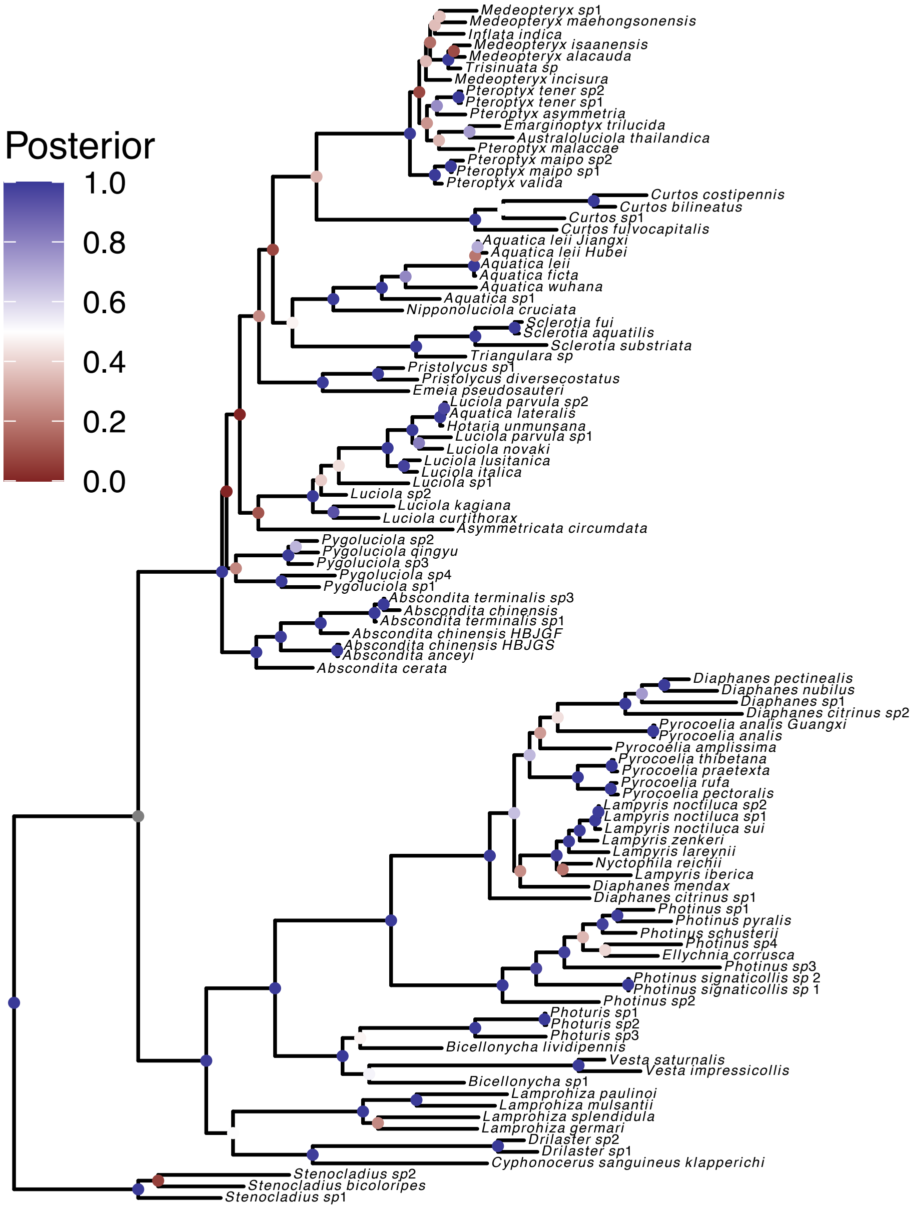

**Figure S14: Maximum a posteriori gene tree for the protein coding gene NADH-6 locus.** Here we show the *maximum a posteriori* (MAP) gene tree, i.e., the gene tree with the highest posterior probability. Circles at internal nodes are colored based on posterior probabilities. The tree was rooted with *Stenocladius* as outgroup.

### S3 Posterior probabilities of specific nodes from single gene analyses

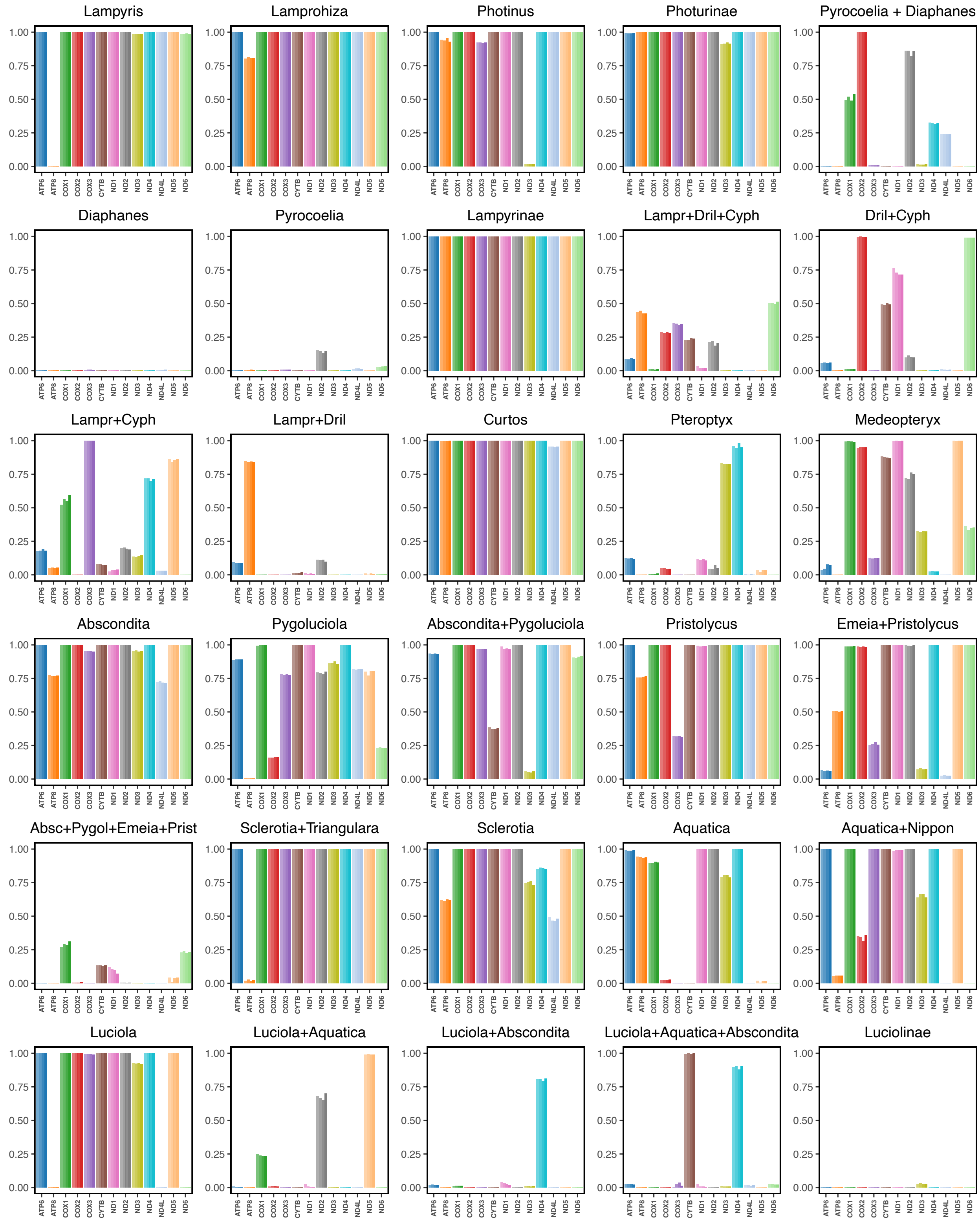

**Figure S15: Posterior probabilities of specific nodes (i.e., clades) obtained from single gene tree analysis under a *uniform* partition model.** Here we show the posterior probabilities of the given clades being monophyletic for 4 replicated MCMC analyses for each of the genes.

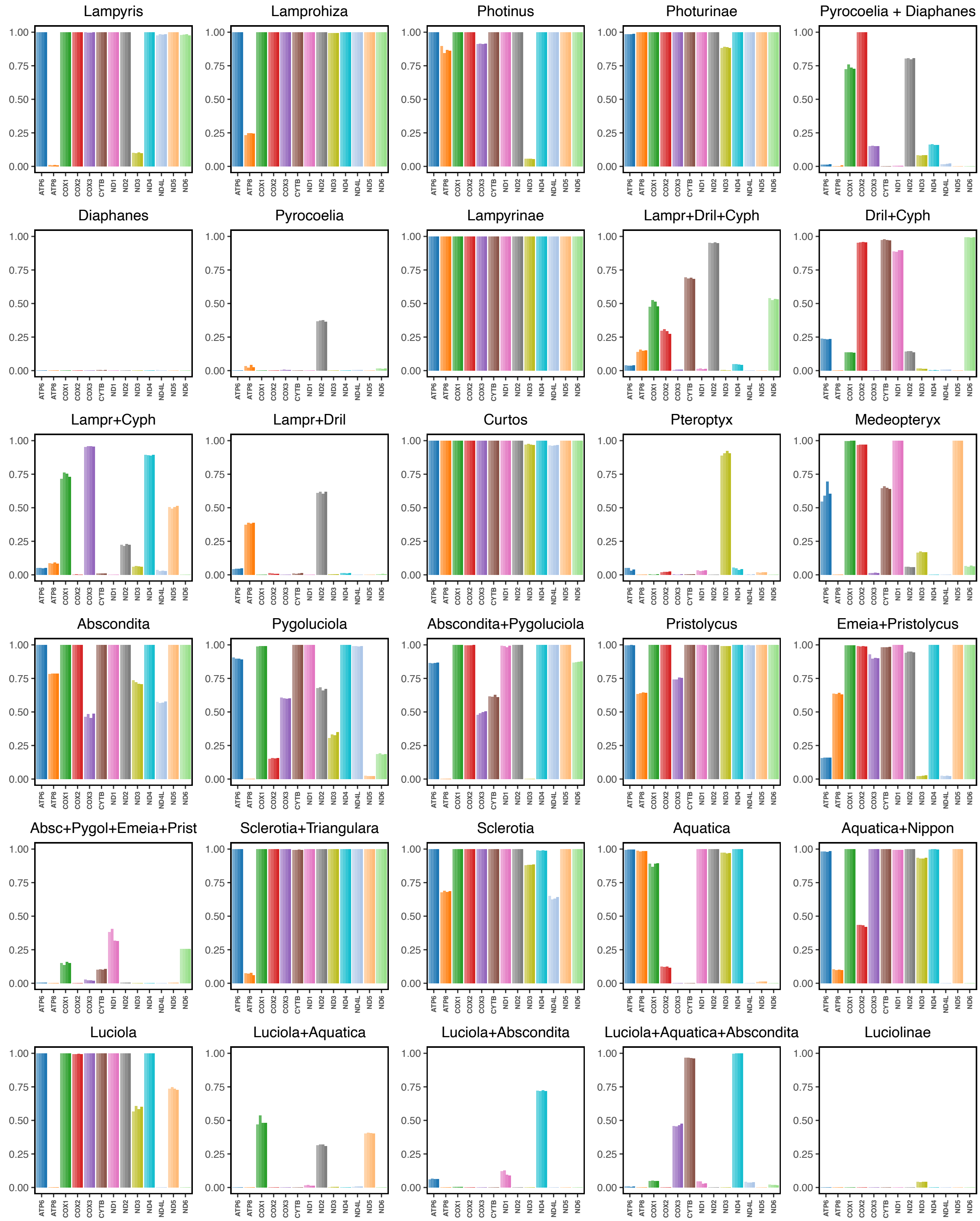

**Figure S16: Posterior probabilities of specific nodes (i.e., clades) obtained from single gene tree analysis under a by *codon* position partition model.** Here we show the posterior probabilities of the given clades being monophyletic for 4 replicated MCMC analyses for each of the genes.

### S4 Unrooted phylogenies using concatenated mitochondrial genomes

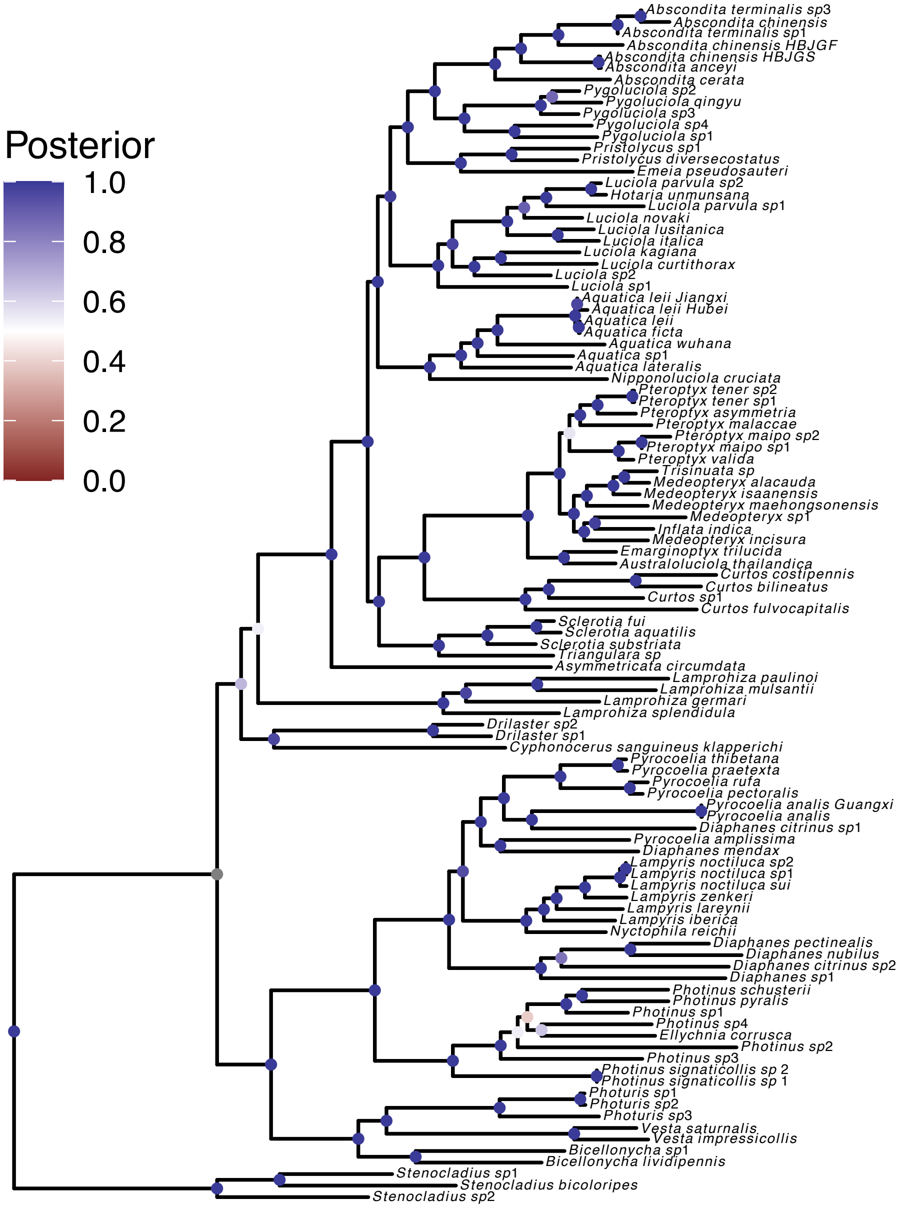

**Figure S17: Maximum a posteriori phylogeny using a concatenation of all 13 protein coding genes plus 2 rRNA genes assuming the *uniform* partitioning scheme.** The *uniform* partitioning scheme assumes that all genes evolve under the same substitution process and substitution rates, i.e., no among-partition rate variation. Here we show the *maximum a posteriori* (MAP) phylogeny, i.e., the species tree with the highest posterior probability. Circles at internal nodes are colored based on posterior probabilities. The tree was rooted with *Stenocladius* as outgroup.

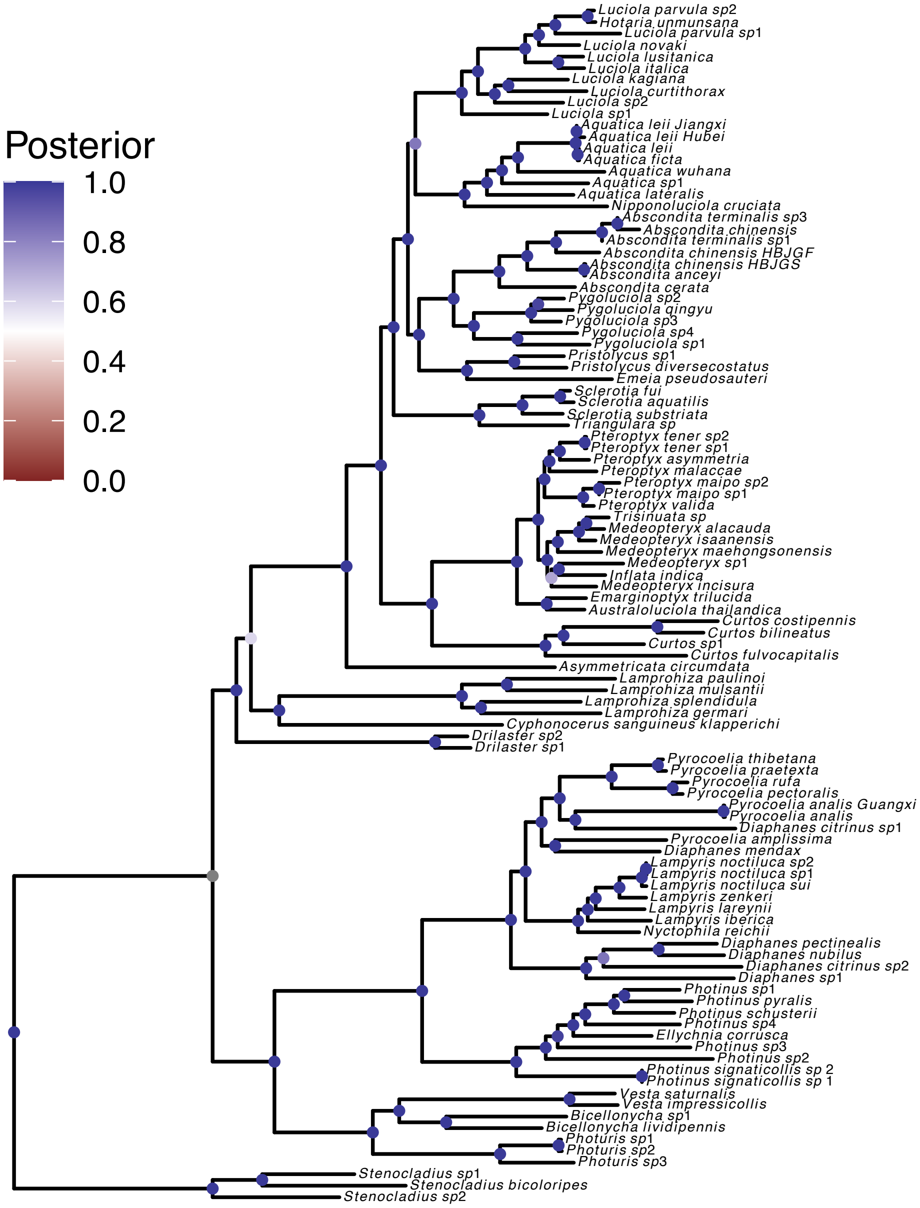

**Figure S18: Maximum a posteriori phylogeny using a concatenation of all 13 protein coding genes plus 2 rRNA genes assuming the by *gene* partitioning scheme.** The by *gene* partitioning scheme assumes that all genes evolve under a separate substitution process and substitution rates, i.e., among-partition rate variation by gene. Here we show the *maximum a posteriori* (MAP) phylogeny, i.e., the species tree with the highest posterior probability. Circles at internal nodes are colored based on posterior probabilities. The tree was rooted with *Stenocladius* as outgroup.

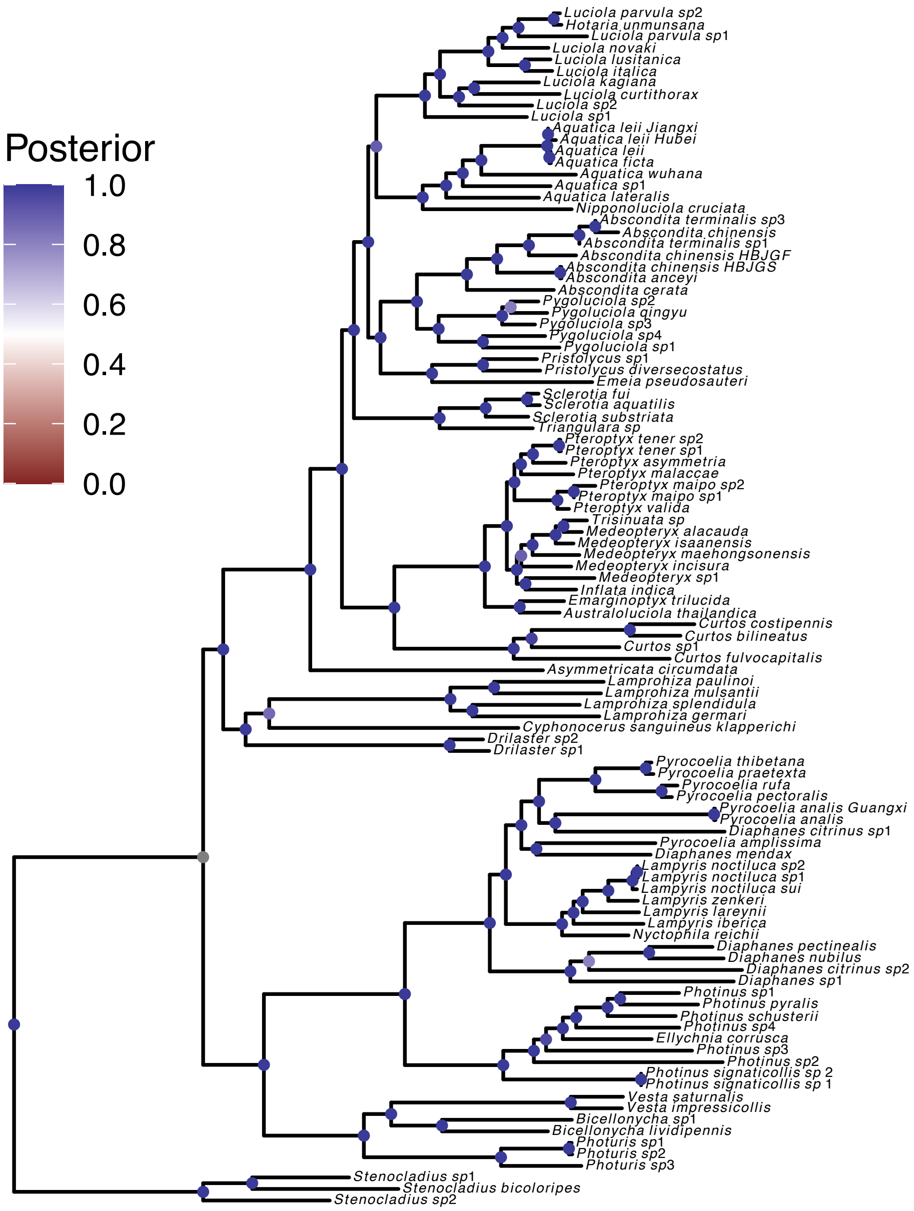

**Figure S19: Maximum a posteriori phylogeny using a concatenation of all 13 protein coding genes plus 2 rRNA genes assuming the by *codon* position partitioning scheme.** The by *codon* position partitioning scheme assumes that all genes evolve under the same substitution process and substitution rates, however, the protein coding genes are split by codon position into three separate data subsets. Additionally, the 2 rRNA genes are placed into their own data subset. Thus, we split the data into 4 data subsets and apply among-partition rate variation. Here we show the *maximum a posteriori* (MAP) phylogeny, i.e., the species tree with the highest posterior probability. Circles at internal nodes are colored based on posterior probabilities. The tree was rooted with *Stenocladius* as outgroup.

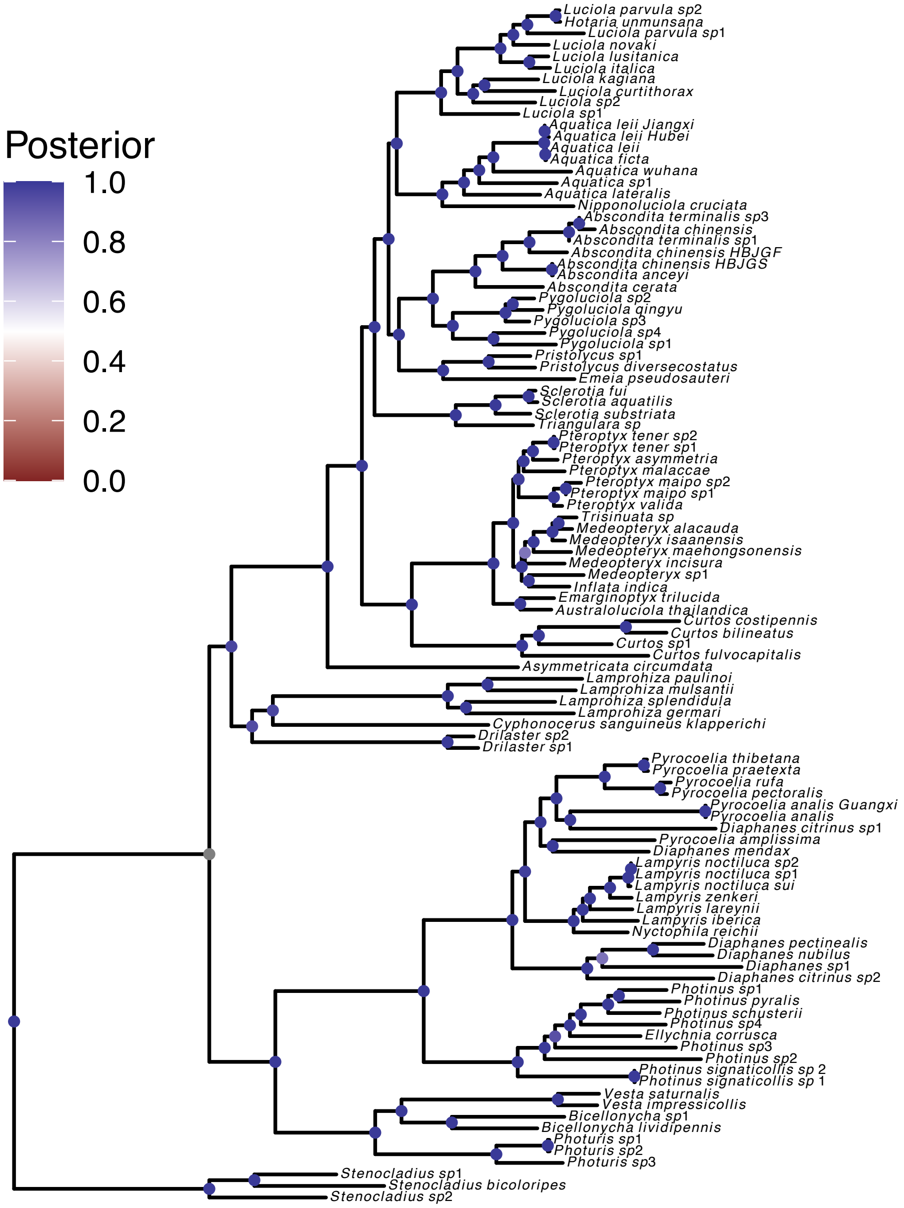

**Figure S20: Maximum a posteriori phylogeny using a concatenation of all 13 protein coding genes plus 2 rRNA genes assuming the by-gene and by-codon position(*combined*) partitioning scheme.** The by-gene and by-codon position (*combined*) partitioning scheme assumes that all genes evolve under a separate substitution process and substitution rates. Furthermore, we split each gene by codon position and place the 2 rRNA genes in their own data subset. Thus, we split the total dataset into 41 data subsets and apply among-partition rate variation. Here we show the *maximum a posteriori* (MAP) phylogeny, i.e., the species tree with the highest posterior probability. Circles at internal nodes are colored based on posterior probabilities. The tree was rooted with *Stenocladius* as outgroup.

### S5 Posterior probabilities of specific nodes from concatenated multilocus analyses

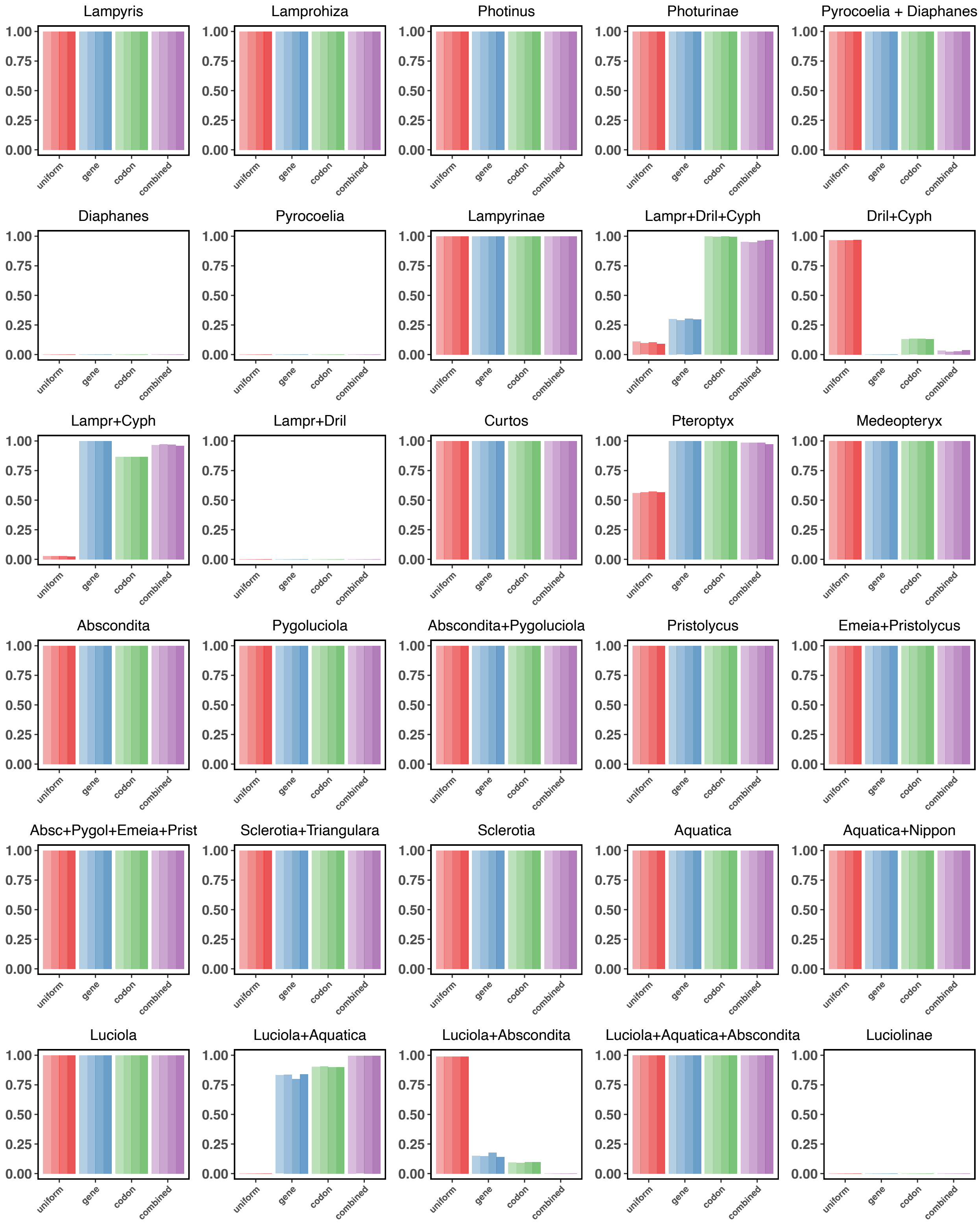

**Figure S21: Posterior probabilities of specific nodes (i.e., clades) obtained from concatenated multi-locus unrooted tree analysis under the four different partition model.** For each partition model we show the posterior probabilities of the given clades being monophyletic for 4 replicated MCMC analyses.

### S6 Time-calibrated phylogenies using concatenated mitochondrial genomes

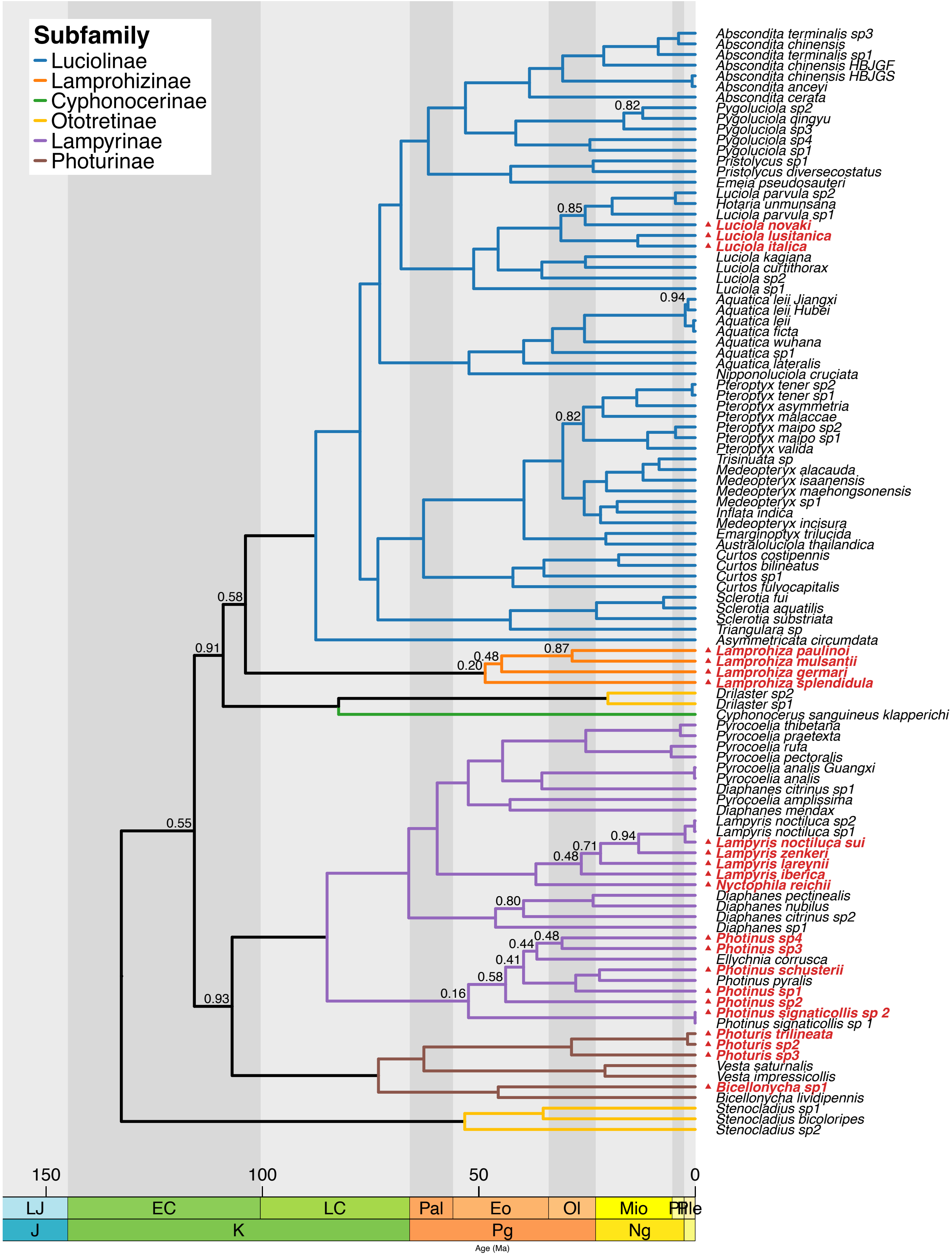

**Figure S22: Maximum a posteriori time-calibrated phylogeny using a concatenation of all 13 protein coding genes plus 2 rRNA genes assuming the *uniform* partitioning scheme.** The *uniform* partitioning scheme assumes that all genes evolve under the same substitution process and substitution rates, i.e., no among-partition rate variation. Here we show the *maximum a posteriori* (MAP) phylogeny, i.e., the species tree with the highest posterior probability. Nodes that have a posterior probability $< 0.95$ are written on the branches above. Branches are colored by subfamily.

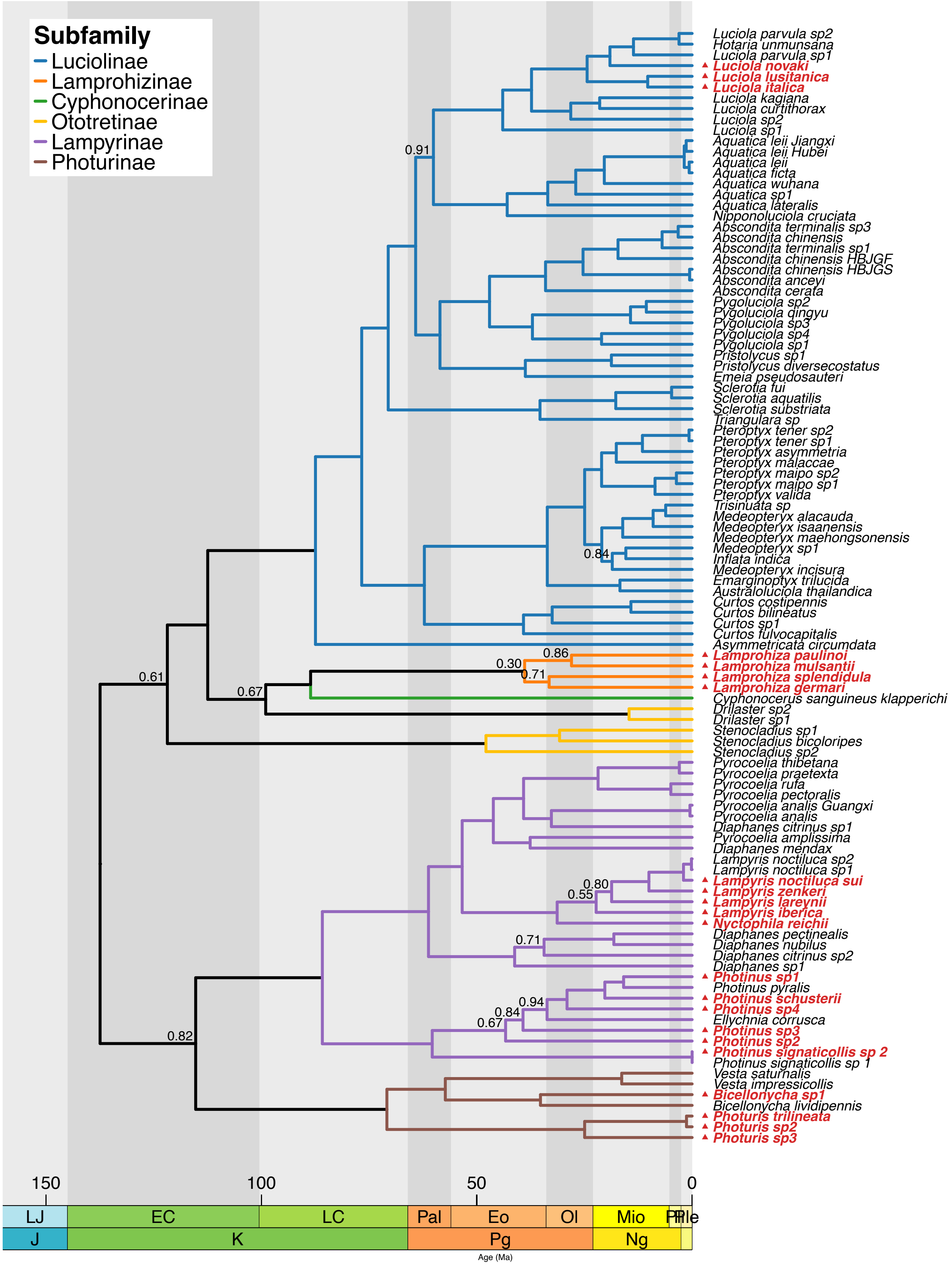

**Figure S23: Maximum a posteriori time-calibrated phylogeny using a concatenation of all 13 protein coding genes plus 2 rRNA genes assuming the by *gene* partitioning scheme.** The by *gene* partitioning scheme assumes that all genes evolve under a separate substitution process and substitution rates, i.e., among-partition rate variation by gene. Here we show the *maximum a posteriori* (MAP) phylogeny, i.e., the species tree with the highest posterior probability. Nodes that have a posterior probability $< 0.95$ are written on the branches above. Branches are colored by subfamily.

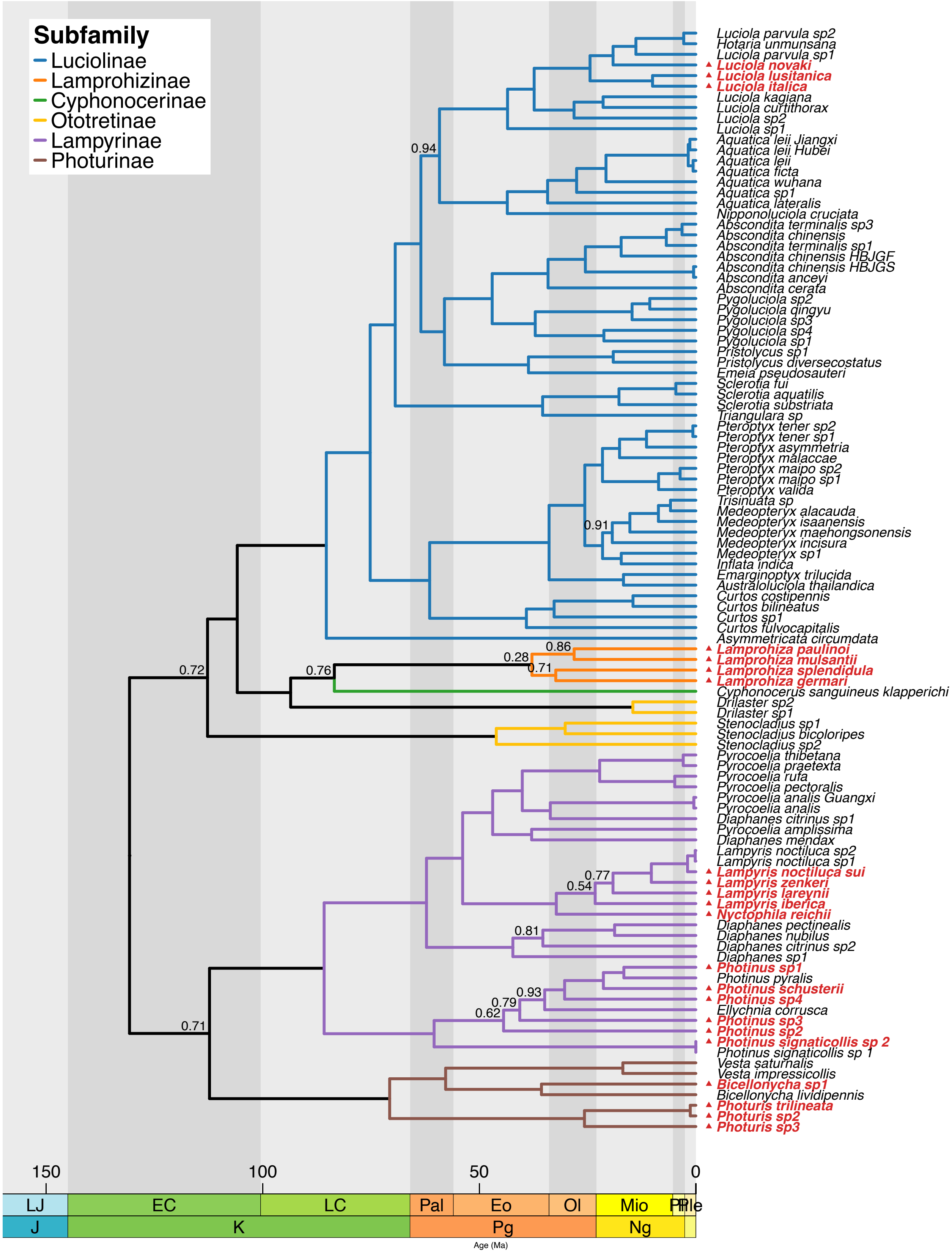

**Figure S24: Maximum a posteriori time-calibrated phylogeny using a concatenation of all 13 protein coding genes plus 2 rRNA genes assuming the by *codon* position partitioning scheme.** The by *codon* position partitioning scheme assumes that all genes evolve under the same substitution process and substitution rates, however, the protein coding genes are split by codon position into three separate data subsets. Additionally, the 2 rRNA genes are placed into their own data subset. Thus, we split the data into 4 data subsets and apply among-partition rate variation. Here we show the *maximum a posteriori* (MAP) phylogeny, i.e., the species tree with the highest posterior probability. Nodes that have a posterior probability $< 0.95$ are written on the branches above. Branches are colored by subfamily.
